# EWS::FLI1-DHX9 interaction promotes Ewing sarcoma sensitivity to DNA topoisomerase 1 poisons by altering R-loop metabolism

**DOI:** 10.1101/2023.05.30.542894

**Authors:** Joaquin Olmedo-Pelayo, Esperanza Granado-Calle, Daniel Delgado-Bellido, Laura Lobo-Selma, Carmen Jordán-Pérez, Ana T. Monteiro-Amaral, Anna C. Ehlers, Shunya Ohmura, Angel M. Carcaboso, Javier Alonso, Isidro Machado, Antonio Llombart-Bosch, Thomas G.P. Grünewald, Fernando Gómez-Herreros, Enrique de Álava

## Abstract

Drug resistance is one of the major factors associated with poor outcome of cancer patients. Treatment of Ewing sarcoma (EwS), an aggressive neoplasm mainly affecting children, adolescents and young adults, is associated with therapy failure and tumor relapse in 30-80% of the cases. Thus, it supports the need to explore the mechanisms modulating drug activity. Here, we describe a novel mechanism of drug sensitivity based on the role of EWS::FLI1 in R-loop metabolism. Our results demonstrate that EWS::FLI1 promotes R-loop formation favoring the interaction between DHX9 and elongating RNA polymerase II. In addition, we discovered that EWS::FLI1 kidnaps DHX9 preventing the resolution of TOP1 poisoning-associated R-loops. Our findings indicate that R-loops accumulation promotes replicative stress, genome instability and cell sensitivity to SN-38. Collectively, these results uncover a novel mechanism behind EwS sensitivity to genotoxic agents, with relevant implications for EwS treatment.

## Introduction

Ewing sarcoma (EwS) is an aggressive mesenchymal malignancy mainly occurring in children, adolescents and young adults. Although 5-year survival for patients with localized disease is ∼70-80%, patients with metastatic disease still present a dismal prognosis with long-term survival rates of ∼30%^1^. In fact, 25-30% of patients present metastasis at diagnosis^2,3^. Despite the collective efforts to implement novel therapeutic approaches, current EwS treatment remains based on the combination of traditional chemotherapy, radiotherapy, and surgery^4^. The prevailing first-line chemotherapy for EwS is the VDC/IE regimen, which includes vincristine, doxorubicin and cyclophosphamide alternating with ifosfamide and etoposide^5,6^. Therapy failure and tumor relapse occur in 30-35% of patients with primary localized disease and 50-80% of the cases with metastasis^4^. Recently, other genotoxic agents (e.g., irinotecan and temozolomide) have been successfully evaluated in clinical trials as effective treatment options in relapsed or advanced refractory EwS^7,8^.

Genetically, EwS is characterized by pathognomonic chromosomal translocations causing gene fusions between TET and ETS gene family members, mainly affecting *EWSR1* and *FLI1* genes (in 85% of the cases)^1,9^. Although EWS::FLI1 is the established main driver of EwS and the frequency of additional mutations is low^10^, the relationship between EWS::FLI1 and the response to genotoxic agents has not been extensively addressed. Strikingly, *Gorthi and colleagues* recently described a novel mechanism involving R-loop formation through EWS::FLI1-mediated increased transcription^11^. They propose that the recruitment of BRCA1 to the elongating transcription machinery would reduce the homologous recombination capacity of EwS cells giving rise to the ‘BRCAness’ phenotype previously associated with EwS tumors.

R-loops are DNA:RNA hybrids generated during transcription when the nascent RNA invades the double-stranded DNA^12^. Despite its physiological roles in many cellular processes, such as DNA replication or gene expression, the accumulation of R-loops is a source of genome instability^13,14^. Different mechanisms finely regulate R-loop levels, including RNA processing factors, topoisomerases, and chromatin remodelers^15^. Additionally, enzymes that degrade the RNA component of R-loops, such as RNAse H1 and RNAse H2^16^, and enzymes with DNA/RNA helicase activity such as DEAD-Box RNA helicases or DEAH helicase DHX9 have been implicated in R-loops metabolism^17,18^.

DHX9 (also known as RNAse helicase A (RHA)) is an NTP-dependent helicase with DNA/RNA helicase activity involved in different cellular processes such as DNA replication, transcription, or RNA processing^19^. The role of DHX9 in R-loop metabolism is not completely understood. Depending on the cellular context, DHX9 can have a negative or a positive effect on R-loop accumulation. On the one hand, the DHX9-mediated unwinding of nascent RNA promotes R-loop formation when splicing is impaired and the interaction of DHX9 and elongating RNAPII is prolonged along gene bodies^20^. On the other hand, DHX9 has been also implicated in the resolution of both physiological and topoisomerase I (TOP1) poisoning-induced R-loops, preventing the generation of genome instability^18,21,22^. Notably, DHX9 interacts with EWS::FLI1 as a transcriptional cofactor enhancing EWS::FLI1-associated aberrant transcription^23,24^. Recently, the small molecule YK-4-279 has shown efficacy in blocking EWS::FLI1-DHX9 binding and, consequently, recovering DHX9 activity^25^.

In this work, we have studied the mechanism underlying R-loop accumulation in EwS cells and its potential implication in the sensitivity to genotoxic agents. We demonstrate that EwS sensitivity to TOP1 poison SN-38, the active molecule of irinotecan, depends on EWS::FLI1-DHX9 interaction. Our findings indicate that the recovery of DHX9 activity enhances R-loop resolution in EwS, reducing drug-associated replicative stress, DNA damage and cytotoxicity.

Overall, these results represent a potential rationale for the association of R-loop accumulation and drug hypersensitivity in EwS.

## Results

### EwS cell lines and tumors are highly sensitive to TOP1 poisons

To elucidate the molecular mechanisms underlying drug sensitivity in EwS, we interrogated the Genomics of Drug Sensitivity in Cancer database^26^. We compared the activity of more than 300 compounds on 15-17 EwS cell lines with respect to more than 600 non-EwS cells. Cell lines bearing EWS::FLI1 fusion protein were significantly more sensitive to drugs commonly used as chemotherapeutic agents, such as PARP inhibitors and DNA damaging agents. Interestingly, we found a marked hypersensitivity of EwS cells to DNA topoisomerase 1 (TOP1) poisons and inhibitors, including irinotecan, topotecan and camptothecin (Fig. 1A; Supplementary Fig. 1A). To validate these observations, we compared the sensitivity of EwS (A4573, A-673, TC-71, EW-7) and non-EwS (U2OS, SK-UT-1, HT-1080, 93T449) cell lines to SN-38, the active metabolite of irinotecan, which confirmed that EwS cell lines are significantly more sensitive toward SN-38 than non-EwS cell lines (Fig. 1B).

**Figure 1.**
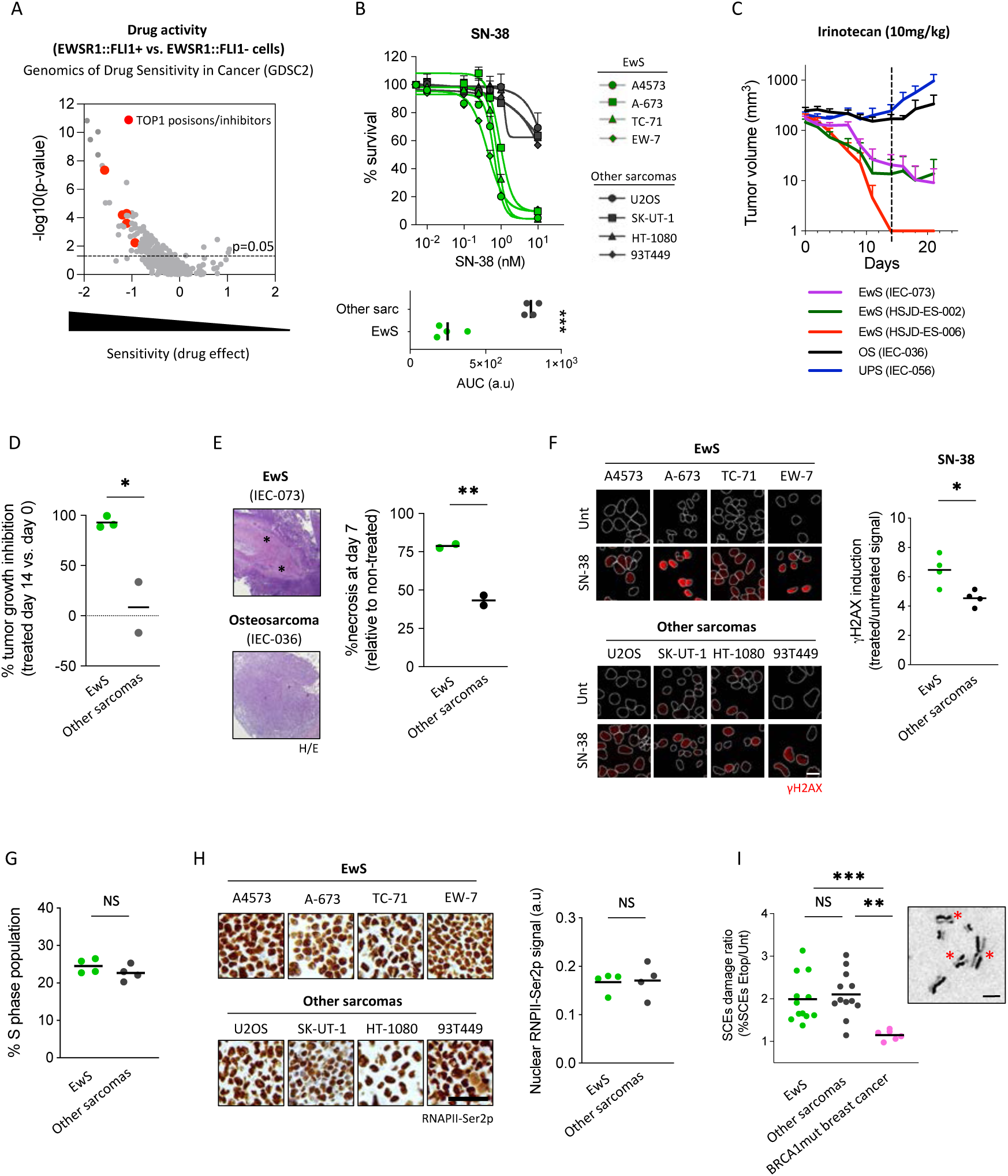
EwS cell lines and tumors are highly sensitive to TOP1 poisons. **(A)** Comparative analysis of drug activity between 15-17 EwS (EWSR1::FLI1) and more than 600 non-EwS cell lines using Genomics of Drug Sensitivity in Cancer website. Each dot is a tested drug. **(B)** Study of the sensitivity to SN-38, comparing EwS (A4573, A-673, TC-71 and EW-7) and non-EwS cell lines (U2OS (osteosarcoma), SK-UT-1 (leiomyosarcoma), HT-1080 (fibrosarcoma) and 93T449 (liposarcoma)). After 72 h of treatment with indicated concentrations, cell viability was analyzed by MTT. *Bottom*, data represent mean of the area under curve of MTT results, n=2 independent experiments. **(C)** Evaluation of tumor growth of PDX models (EwS: IEC-073, HSJD-ES-002, HSJD-ES-006; OS, osteosarcoma, IEC-036; UPS, undifferentiated pleomorphic sarcoma, IEC-056) over 21 days following the beginning of irinotecan treatment. Dotted line indicates the end of treatment. **(D)** Evaluation of tumor growth inhibition of PDX models at the end of irinotecan treatment. Data represent mean of the percentage of tumor growth inhibition. **(E)** Evaluation of tumor necrosis at day 7 of treatment. *Left*, representative hematoxylin/eosin images. Asterisks indicate necrotic areas. *Right*, data represent mean of the percentage of tumoral necrosis. **(F)** Analysis of drug-induced DSBs comparing EwS (A4573, A-673, TC-71 and EW-7) and non-EwS cell lines (U2OS, SK-UT-1, HT-1080 and 93T449) by γH2AX IF. Cells were treated with 5 µM SN-38 (30 min). *Left*, representative images. DAPI counterstain. Scale bar, 20 µm. *Right*, data represent mean of nuclear γH2AX intensity, n≥3 independent experiments. **(G**) Analysis of the fraction of cells in S phase of EwS and non-EwS cell lines by FACS (propidium iodide staining). **(H)** Evaluation of transcriptional activity of EwS and non-EwS cell lines by RNAPII-Ser2p ICC in paraffin-embedded pellets. *Left*, representative images. Scale bar, 50 µm. *Right*, data represent mean of nuclear RNAPII-Ser2p signal. **(I)** Study of HR efficiency by SCE assay in EwS, non-EwS and BRCA mutated breast cancer cell lines (MDA-MB-436 and HCC-1937). *Left*, data represent mean of SCEs induction upon 2.5 µM etoposide treatment (30 min), n=3 independent experiments. *Right*, representative image. Scale bar, 5 µm. Asterisks indicate SCEs events. Statistical significance was determined by t-test (NS, not significant; *P<0.05; **P<0.01; ***P<0.001).

Next, we studied irinotecan sensitivity *in vivo* using EwS (IEC-073, HSJD-ES-002, HSJD-ES-006) and non-EwS (IEC-036, osteosarcoma, OS; IEC-056, undifferentiated pleomorphic sarcoma, UPS) patient-derived xenografts (PDXs) models. Tumors were subcutaneously implanted and treated as previously described^27^. Evaluation of tumor volume during 21 days after the beginning of the treatment showed a higher response of EwS compared to non-EwS models (Fig. 1C), which is concordant with the potent activity of irinotecan in Ewing sarcoma PDXs^28^. When calculating tumor growth inhibition compared to tumor volume at the end of treatment (day 14), we found that growth inhibition was significantly higher in EwS PDX models (92.83%) compared to OS (33.67%) and UPS (-17.02%) (Fig. 1D). Indeed, 95% (19/20) of EwS tumors showed a complete response to irinotecan while most of OS and UPS tumors showed a stable response or tumor progression disease (Supplementary Fig. 1B). Tumoral response was also assessed by evaluating the treatment-induced necrosis at day 7, which was significantly higher in EwS PDXs (78.75%) compared to OS (40%) and UPS (46.5%) models (Fig. 1E). Finally, evaluation of animal event-free survival showed that EwS PDX models survived more than 60 days without evidence of tumor growth, while OS and UPS animals were sacrificed upon achieving endpoint (tumor volume around 1200 mm^3^) at 32.5 and 26.5 days after the start of the treatment, respectively (Supplementary Fig. 1C).

Altogether, these results demonstrated that, both *in vitro* and *in vivo*, EwS are significantly more sensitive to TOP1 poisons than other tested sarcoma models.

### EwS sensitivity to TOP1 poisons is independent of proliferation, transcription, and recombination

TOP1-poisoning-induced single-strand breaks are readily converted into double-strand breaks (DSBs), the most cytotoxic form of DNA damage, mainly through collision with replication machinery^29^. Consequently, we analyzed whether the hypersensitivity of EwS cell lines to TOP1 poisons could be associated with a stronger induction of DSBs. Indeed, evaluation of H2AX serine 139 phosphorylation (hereafter γH2AX), a surrogate marker of DSBs^30^, indicated that SN-38 treatment induced higher levels of DSBs in EwS in comparison to non-EwS cells (Fig. 1F). Genotoxic agents present different mechanisms for DSB induction; nevertheless, there are common cellular factors that modulate the formation of these lesions, such as proliferation or transcription. Therefore, differences in the activity of these processes could vary in DSB induction and drug cytotoxicity. To analyze potential differences in proliferation, we first measured the percentage of cells in S-phase by flow cytometry, which yielded no significant differences between EwS and non-EwS cells (Fig. 1G). Similarly, proliferation rates were analyzed by Ki-67 immunocytochemistry (ICC) in paraffin-embedded cell lines pellets showing no significant difference in Ki-67+ staining between both groups (Supplementary Fig. 1D). MRC-5, a non-tumoral fibroblast cell line with low proliferation rate, was included as a control. Next, we immunohistochemically evaluated global transcription by detecting Serine 2 phosphorylation of the C-terminal domain of RNA polymerase II (hereafter RNAPII-Ser2p), a marker of active transcription. Results showed similar levels of RNAPII-Ser2p in EwS and non-EwS cells (Fig. 1H). Additionally, we did not observe differences in nascent RNA levels measured by uridine analogue, 5-ethynyluridine (EU) incorporation (Supplementary Fig. 1E). These results suggest that EwS cells have similar levels of proliferation, DNA replication or transcription to non-EwS cells.

Finally, we reasoned that ‘BRCAness’ phenotype previously proposed in association with EwS could be the cause of DSB accumulation upon treatment with DNA-damaging agents. To study the putative ‘BRCAness’ phenotype in EwS cells, we analyzed sister chromatid exchanges (SCEs), a well-established hallmark of homologous recombination, upon DSB induction with a low dose of etoposide. However, no significant differences were observed between EwS and non-EwS cells (Fig. 1I; Supplementary Fig. 1F). Notably, in all cases, induction of SCEs was significantly higher compared to two breast cancer cell lines (BRCA1 mutated) (Fig. 1I; Supplementary Fig 1F). Taken together, these results indicated that the hypersensitivity of EwS cells to TOP1 poisons cannot be ascribed to higher proliferation, DNA replication or transcription nor to lower HR efficiency compared to other tumoral cells.

### EWS::FLI1 impairs drug-induced R-loop resolution promoting genome instability and cytotoxicity

Since TET-ETS fusions are the main drivers of EwS, we studied the contribution of EWS::FLI1 to the hypersensitivity to TOP1 poisons. Herein, efforts were focused on the pre-established A-673/TR/shEF model (hereafter shA673), carrying a doxycycline (DOX) inducible shRNA against EWSR1::FLI1^31^. shA673 cells exposed to DOX exhibited decreased EWS::FLI1 both at mRNA and protein levels (Fig. 2A; Supplementary Fig. 2A), leading to an upregulation of LOX (EWS::FLI1 repressed target)^32^ and a gradual accumulation of cells in G1 phase of the cell cycle, as previously described (Supplementary Fig. 2A, 2B). Given that changes in cell proliferation could affect drug activity, experiments were carried out at 24 hours of DOX incubation. At this time point, no significant differences were neither observed in cell cycle nor replication activity as measured by thymidine analogue, 5-Ehylnyl-2’-deoxyuridine (EdU) incorporation (Supplementary Fig. 2B, 2C). To analyze the contribution of EWS::FLI1 to SN-38 cytotoxicity, shA673 cells were incubated with DOX for 24 h before treatment with SN-38. Notably, EWS::FLI1 depletion reduced SN-38-associated cleavage of PARP1 and Casp3, suggesting that downregulation of EWS::FLI1 reduces drug-induced apoptosis (Fig. 2B). In agreement, EWS::FLI1 knockdown significantly reduced the percentage of drug-induced AnnexinV+ cells (Fig. 2C). Finally, in accordance with these results, downregulation of EWS::FLI1 significantly increased cell survival to SN-38 (IC50 from 0.07 to 0.2 μM) (Fig. 2D), demonstrating that EWS::FLI1 promotes sensitivity to TOP1 poisons in EwS cells. Next, to elucidate whether EWS::FLI1 expression modulates SN-38-induced DSBs, we analyzed γH2AX levels upon DOX incubation. Strikingly, downregulation of EWS::FLI1 significantly reduced the levels of SN-38-induced DNA damage (Fig. 2E). Since γH2AX is a surrogate DSBs marker, we directly measured chromosomal breaks in spread preparations. In accordance with our previous results, we observed that pre-treatment with DOX significantly reduced drug-induced chromosomal breaks (Fig. 2F).

**Figure 2.**
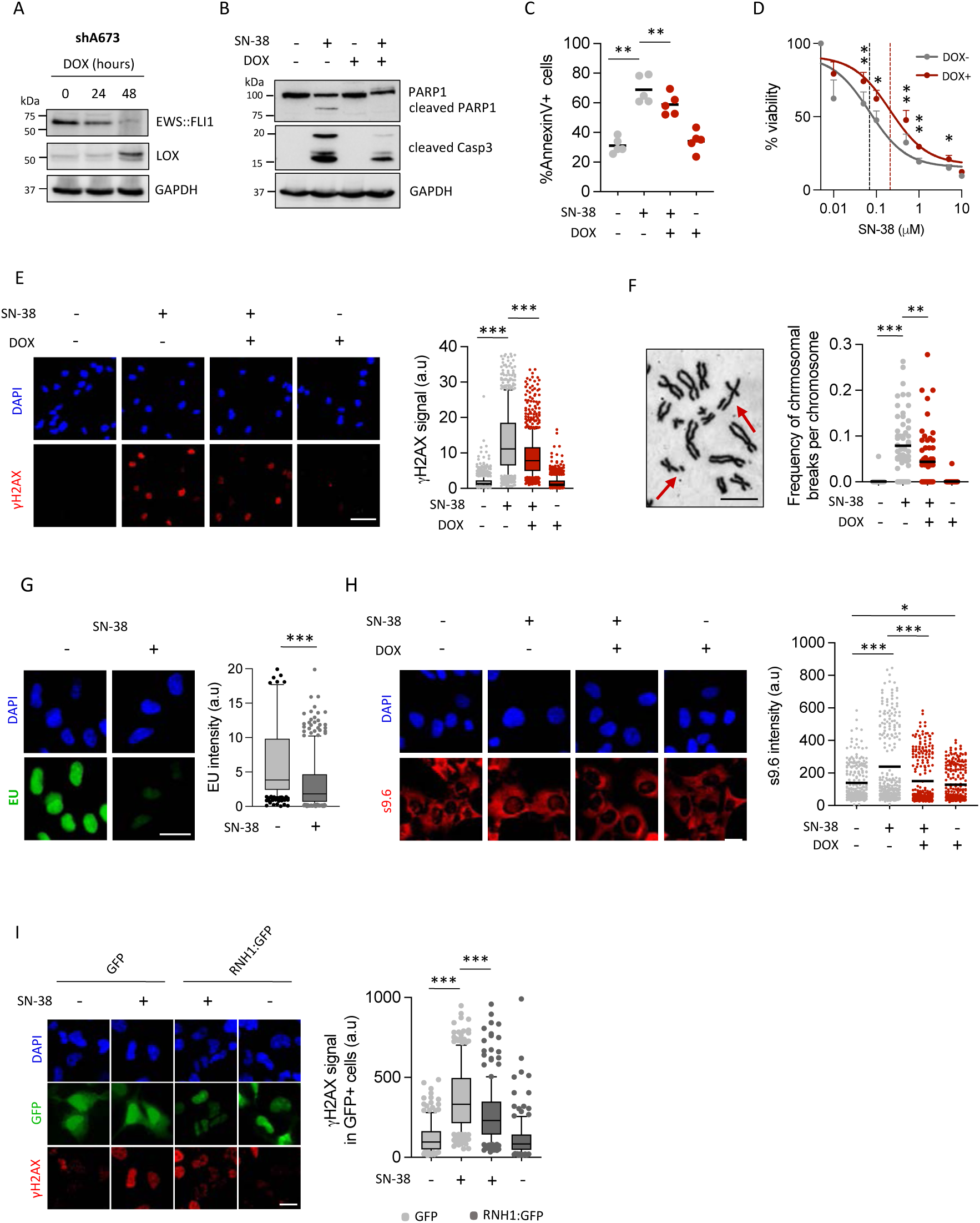
EWS::FLI1 impairs drug-induced R-loop resolution promoting genome instability and cytotoxicity. **(A)** EWS::FLI1 and LOX protein levels by WB in shA673 cells after DOX incubation. Loading control: GAPDH. Molecular weight in kDa. **(B)** SN-38 induced apoptosis upon EWS::FLI1 knockdown by PARP1 and CASP3 WB. Cells were pre-incubated with DOX before 5 µM SN-38 treatment (3 h). After washout, cells were cultured in drug-DOX-free medium for 24 h. Loading control: GAPDH. Molecular weight in kDa. **(C)** Similar to (B) by AnnexinV FACS (24 h after washout). Data represent mean of the percentage of AnnexinV+ cells, n=5 independent experiments. **(D)** Similar to (B). Cells were treated with indicated concentrations of SN-38 (3 h). After treatment, cells were cultured in drug-DOX-free medium for 48 h before MTT. Data represent mean (±SEM) of the percentage of survival, n=5 independent experiments. Dotted lines indicate IC50. **(E)** SN-38 induced γH2AX upon EWS::FLI1 knockdown by IF. Cells were pre-incubated with DOX and treated with 5 µM SN-38 (30 min). *Left*, representative images. DAPI counterstain. Scale bar, 50 µm. *Right*, data represent mean (±10-90 percentile) of nuclear γH2AX intensity, n=5 independent experiments. **(F)** SN-38 induced chromosomal breaks upon EWS::FLI1 downregulation. After DOX incubation, cells were treated with 2.5 µM SN-38 (30 min). *Left,* representative image. Scale bar, 10 µm. Arrows indicate chromosomal breaks. *Right,* data represent the mean of the frequency of chromosomal breaks per chromosome, n=3 independent experiments. **(G)** Evaluation of transcription by EU incorporation. shA673 cells were treated with 5 µM SN-38 (30 min). *Left,* representative images. DAPI counterstain. Scale bar, 20 µm. *Right*, data represent mean (±10-90 percentile) of EU signal, n=4 independent experiments. **(H)** SN-38 induced R-loops upon EWS::FLI1 knockdown by s9.6 IF. Cells were pre-incubated with DOX and treated with 5 µM SN-38 (30 min). *Left*, representative images. DAPI counterstain. Scale bar, 20 µm. *Right*, data represent mean of nuclear s9.6 intensity, n=3 independent experiments. **(I)** Effect of RNH1 overexpression on SN-38-induced DSBs. Cells were transfected with RNH1:GFP or control plasmids 48 h before 5 µM SN-38 treatment (30 min). *Left,* representative images. DAPI counterstain. Scale bar, 20 µm. *Right.* data represent the mean (±10-90 percentile) of nuclear γH2AX intensity in GFP+ cells, n=3 experiments. P value was determined by t-test (*P<0.05; **P<0.01; ***P<0.001).

Considering that TOP1 poisoning was previously associated with the accumulation of R-loops^33,34^, a prominent source of DNA damage and genome instability, we wondered whether EWS::FLI1 contributes to R-loop metabolism. As previously described, TOP1 poisoning caused a significant decrease in transcription, measured by EU incorporation (Fig. 2G). To analyze the possible accumulation of R-loops upon TOP1 poisoning, we quantified R-loops by immunofluorescence microscopy using s9.6, an R-loop specific antibody. Interestingly, SN-38 treatment induced a significant accumulation of nuclear R-loops. Notably, downregulation of EWS::FLI1 in shA673 cells resulted in a decrease in the levels of SN-38-induced R-loops levels (Fig. 2H), indicating that EWS::FLI1 is involved in the metabolism of drug-induced R-loops. More importantly, R-loops depletion by overexpression of RNH1 in A-673 cells significantly reduced SN38-induced DSBs (Fig. 2I & Supplementary Fig. 2D), demonstrating the contribution of SN38-induced R-loops to the generation of genome instability in EwS cells.

### Loss of EWS::FLI1-DHX9 interaction prevents SN-38-induced R-loops accumulation and genome instability

Given the previously described physical interaction of EWS::FLI1 with DHX9 and the role of DHX9 in the metabolism of R-loops, we hypothesized that EWS::FLI1 could interfere with DHX9 activity preventing the resolution of the hybrids. Thus, we evaluated whether DHX9 levels affect SN-38-induced DNA damage, and cytotoxicity in EwS cells. First, we downregulated DHX9 in A-673 cells by siRNA, resulting in a significant increase of drug-associated γH2AX levels (Fig. 3A, Supplementary Fig. 2E). By contrast, overexpression of DHX9 was associated with a reduction of SN-38-induced DSBs and cytotoxicity (Fig. 3B, 3C and Supplementary Fig 2F), suggesting that DHX9 prevents SN-38-induced DSBs and cell death.

**Figure 3.**
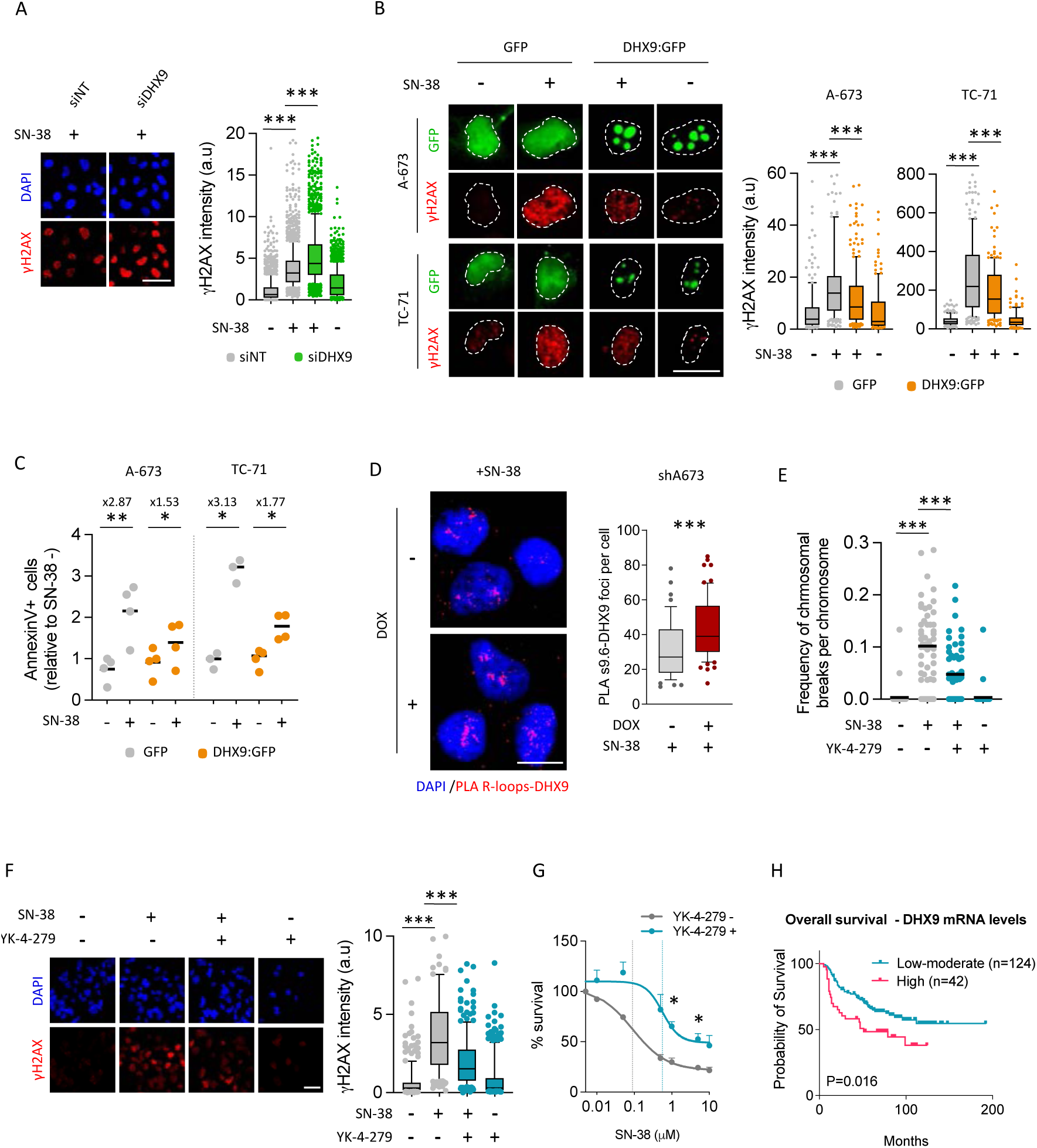
Loss of EWS::FLI1-DHX9 interaction alleviates R-loops accumulation promoting drug resistance. **(A)** Evaluation of the effect of DHX9 knockdown in SN-38 associated DSBs by γH2AX IF. After 48 h of knockdown, A-673 cells were treated with 5 µM SN-38 (30 min). *Left*, representative images. DAPI counterstain. Scale bar, 50 µm. *Right*, data represent the mean (±10-90 percentile) of nuclear γH2AX signal, n=3 independent experiments. **(B)** Evaluation of the effect of DHX9 overexpression on SN-38-induced DSBs by γH2AX IF. A-673 and TC-71 overexpressing DHX9:GFP or control plasmid were treated with 5 µM SN-38 (30 min). *Left*, representative images. DAPI counterstain. Scale bar, 20 µm. *Right*, data represent mean (±10-90 percentile) of nuclear γH2AX signal of GFP+ cells, n≥3 independent experiments. **(C)** Effect of DHX9 overexpression on SN-38-induced apoptosis by AnnexinV FACS. Cells overexpressing DHX9:GFP or control plasmid were treated with 5 µM SN-38 (3 h). After washout, cells were maintained in drug-free medium for 24 h. Data represent mean of the percentage of AnnexinV+ between GFP+ cells, n≥3 independent experiments. **(D)** Determination of R-loops-DHX9 interactions by PLA in shA673 cells upon DOX incubation and 5 µM SN-38 treatment (30 min). *Left*, representative images. DAPI counterstain. Scale bar, 20 µm. *Right*, data represent mean (±10-90 percentile) of PLA foci, n=3 independent experiments. **(E)** Effect of YK-4-279 on SN-38-induced chromosomal breaks. A-673 cells were pre-incubated with 75 µM YK-4-279 (1.5 h) and treated with 2.5 µM SN-38 (30 min). Data represent mean of chromosomal breaks per chromosome, n=3 independent experiments. **(F)** Analysis of the effect of YK-4-279 on SN-38-induced DSBs by γH2AX IF. A-673 cells were pre-incubated with 75 µM YK-4-279 (1.5 h) and treated with 5 µM SN-38 (30 min). *Left*, representative images. DAPI counterstain. Scale bar, 50 µm. *Right*, data represent mean (±10-90 percentile) of nuclear γH2AX intensity, n=3 independent experiments. **(G)** Study of the effect of YK-4-279 on cell survival upon SN-38 exposure. A-673 cells were pre-incubated with 75 µM YK-4-279 (1.5 h) and treated with indicated concentrations of SN-38 (3 h). After treatment, cells were cultured in drug-DOX-free medium for 48 h previous MTT assay. Data represent mean (±SEM) of the percentage of survival, n=3 independent experiments. Dotted lines indicate IC50. Statistical significance was determined by t-test (*P<0.05; **P<0.01; ***P<0.001). **(H)** Kaplan-Meier showing overall survival of 166 EwS patient samples stratified by DHX9 mRNA levels. Statistical significance was determined by Mantel-Cox test.

Next, we explored the role of EWS::FLI1-DHX9 interaction in drug-induced R-loop metabolism. Based on the previous data, we analyzed whether EWS::FLI1 prevents the binding of DHX9 to the R-loops by proximity ligation assay (PLA) using s9.6 and anti-DHX9 antibodies. Results indicated that downregulation of EWS::FLI1 significantly increased the interaction between DHX9 and R-loops (Fig. 3D). To confirm that EWS::FLI1-DHX9 interaction is a source of drug-associated genome instability and cytotoxicity in EwS cells, we used the YK-4-279 inhibitor. Strikingly, pre-treatment of A-673 cells with YK-4-279 significantly reduced SN-38-induced γH2AX and chromosomal breaks (Fig. 3E, 3F). In addition, the loss of EWS::FLI1-DHX9 interaction reduced SN-38 sensitivity (Fig. 3G).

Due to the relevance of DHX9 activity in the resolution of R-loops and the modulation of EwS sensitivity to DNA-damaging agents, we analyzed the relationship between DHX9 levels and EwS patient outcome. Exploration of a cohort including 166 EwS patients^35^ indicated that high levels of DHX9 were associated with reduced overall survival (P = 0.016) (Fig. 3H). In addition, we observed a significant association between high levels of DHX9 expression and a reduced event-free survival^36^ (P = 0.0446) (Supplementary Fig. 2G).

Collectively, these findings indicated that EWS::FLI1-DHX9 interaction prevents SN-38-induced R-loops resolution promoting its accumulation and the induction of genome instability, which may have prognostic relevance.

### SN-38-induced R-loops strengthen replicative stress in EwS

Replicative stress is an important source of R-loop-associated genome instability in cycling cells^13^. Notably, overexpression of RNH1 in A-673 cells significantly reduced the levels of SN-38-induced CHK1 ser345 phosphorylation (pCHK1), a replicative stress marker, indicating that R-loops may mediate replicative stress induced by TOP1 poisons (Fig. 4A). Notably, in agreement with the observed decrease of drug-induced R-loops (Fig. 2H), knockdown of EWS::FLI1 strongly reduced SN-38-induced pCHK1 (Fig. 4B). Similarly, downregulation of EWS::FLI1 was associated with a significant decrease of RPA2 ser4 and ser8 phosphorylation (pRPA2), a different replicative stress marker (Supplementary Fig 3A). To confirm the origin of replicative stress we directly monitored DNA replication by DNA fiber technique measuring incorporation of CldU, followed by IdU, in the presence of SN-38. As previously shown, SN-38 hindered replication progression in shA673 cells (Fig. 4C). Strikingly, EWS::FLI1 depletion prevented this defect. Finally, we compared the levels of drug-associated replicative stress of EwS and non-EwS cells. Results indicated that EwS cells accumulate higher levels of pCHK1 than non-EwS upon SN-38 exposure (Fig 4D).

**Figure 4.**
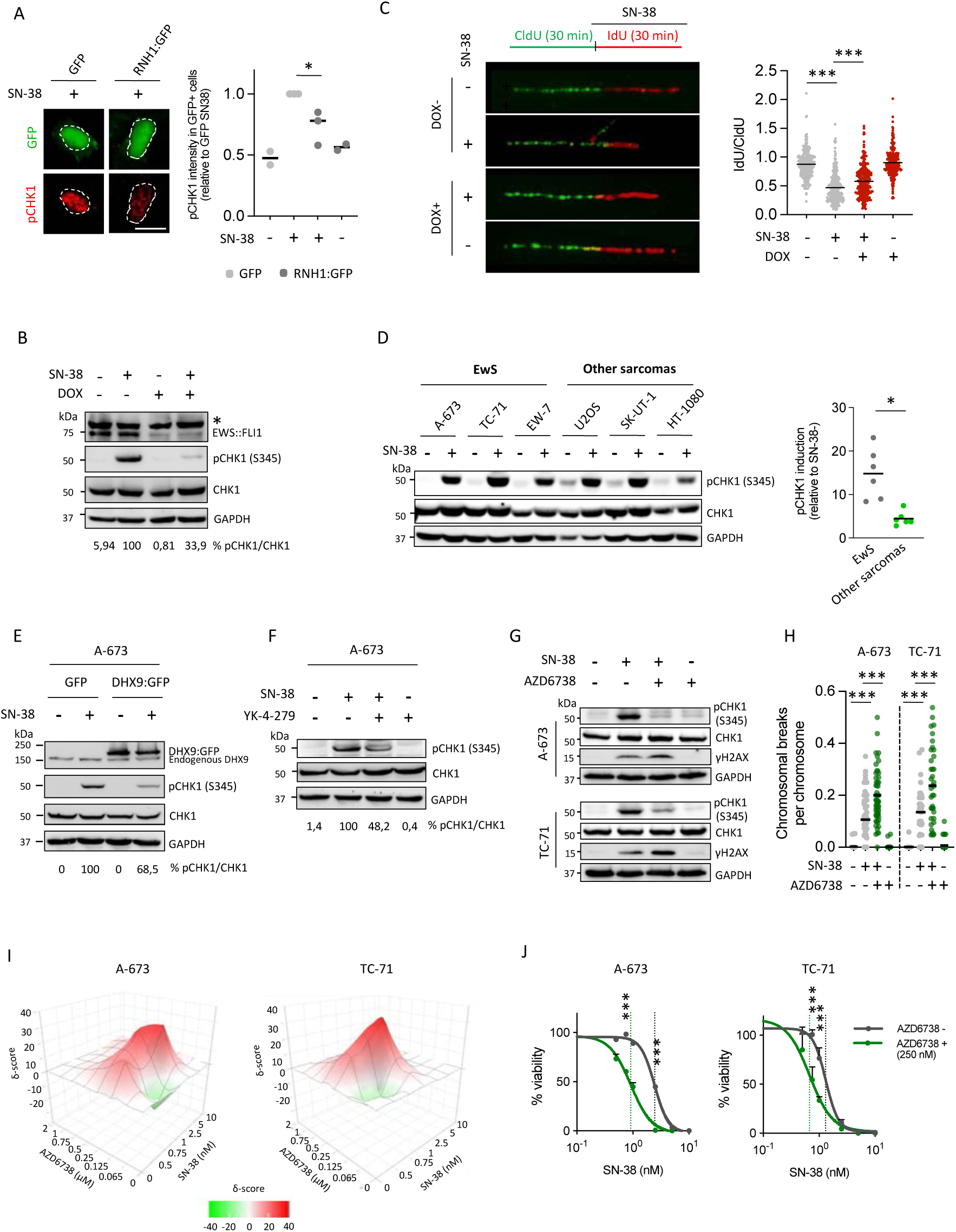
EWS::FLI1-DHX9 associated R-loop accumulation is a source of replicative stress. **(A)** Analysis of the effect of RNH1 overexpression in SN-38-induced replicative stress by pCHK1 (Ser345) IF. Cells were treated with 5 µM SN-38 (30 min). *Left*, representative images. DAPI counterstain. Scale bar, 50 µm. *Right,* data represent mean of pCHK1 intensity in GFP+ cells, n≥2 independent experiments. **(B)** Effect of EWS::FLI1 knockdown in SN-38-induced replicative stress by pCHK1 (Ser345) WB. shA673 cells were pre-incubated with DOX and treated with 5 µM SN-38 (30 min). *Bottom*, quantification of CHK1 phosphorylation (pCHK1/CHK1 band signal). Data are the mean of 2 independent experiments (relative to SN-38). Loading control: GAPDH. Molecular weight in kDa. Asterisk indicates non-specific bands. **(C)** Similar to (B), by DNA fibers assay. Cells were incubated with Cidu (30 min) and IdU + 5 µM SN-38 (30 min). *Left*, representative images. *Right*, mean of IdU/CidU fiber length ratio. Data are the mean of 2 independent experiments (300 fibers per condition). **(D)** Evaluation of SN-38-induced pCHK1 between EwS and non-EwS cells lines. Left, representative immunoblots. Details as in (B)*. Right,* data represent mean of pCHK1 induction, n=2 independent experiments. **(E)** Effect of DHX9 overexpression on SN-38-induced replicative stress by pCHK1 (Ser345) WB. A-673 cells overexpressing DHX9:GFP or control plasmid were treated with 5 µM SN-38 (30 min). *Bottom*, quantification of CHK1 phosphorylation (pCHK1/CHK1 band signal). Data are the mean of 2 independent experiments (relative to SN-38). Other details as in (B). **(F)** Effect of YK-4-279 treatment on SN-38-induced replicative stress by pCHK1 (Ser345) WB. Cells were pre-incubated with 75 µM YK-4-279 (1.5 h) and treated with 5 µM SN-38 (30 min). Other details as in (B). **(G)** Evaluation of the effect of ATRi in SN-38 induced pCHK1 and γH2AX by WB. A-673 and TC-71 cells were pre-incubated with 10 µM AZD6738 for 24 h and treated with 5 µM SN-38 (30 min). Other details as in (B). **(H)** Analysis of the effect of ATRi in SN-38 induced chromosomal breaks. Cells were pre-incubated with 10 µM AZD6738 for 24 h and treated with 2.5 µM SN-38 (A-673) or 1.25 µM SN-38 (TC-71) for 30 min. Data represent mean of the frequency of chromosomal breaks per chromosome, n≥2 independent experiments. Statistical significance was determined by t-test (*P<0.05; **P<0.01; ***P<0.001). **(I)** Bliss synergy plots representing synergistic effect of AZD6738 and SN-38 combination. **(J)** Study of the effect of ATRi on cell survival upon SN-38 exposure by MTT assay. Cells were incubated with AZD6738 (2.5 nM, 24 h) previous to the treatment with indicated concentrations of SN-38 (72 h). Data represent mean (±SEM) of the percentage of survival, n=3 independent experiments. Dotted lines indicate IC50. Statistical significance was determined by two way-anova (***P<0.001).

To evaluate whether EWS::FLI1 promotes replicative stress through the interaction with DHX9, we first measured pCHK1 levels in A-673 and TC-71 upon DHX9 overexpression. Notably, high levels of DHX9 reduced pCHK1 levels induced by SN-38 (Fig. 4E; Supplementary Fig. 3B). More importantly, pre-incubation of A-673 cells with YK-4-279 inhibitor also reduced drug-associated pCHK1 levels (Fig. 4E), confirming that EWS::FLI1-DHX9 interaction promotes replicative stress in EwS cells upon TOP1 poisoning. Taken together, these data indicate that SN-38 induces high levels of replicative stress in EwS cells through the accumulation of R-loops.

Next, we focused on ataxia telangiectasia mutated and rad3 related (ATR), a kinase activated in response to replicative stress that blocks replication fork progression to prevent the generation of DNA damage^37^. ATR inhibition with AZD6738 (ceralasetib), efficiently blocked ATR signaling pathway, reducing pCHK1 levels (Fig. 4G). Interestingly, ATR inhibition increased SN-38-induced γH2AX levels in A-673 and TC-71 cell lines (Fig. 4G). Since ATR phosphorylates H2AX in the context of replicative stress^38^, we directly measured chromosomal breaks in spread preparations. In accordance with our previous result, ATR inhibition increased the levels of SN-38-induced chromosomal breaks in A-673 and TC-71 cells (Fig. 4H), indicating that blockage of replicative stress signaling could enhance SN-38-induced genome instability in EwS cells.

Finally, we analyzed the potential synergistic effects of ATR inhibition and TOP1 poisoning. To this end, cells were pretreated with AZD6738 for 24 h before SN-38 treatment. Results indicated a synergistic effect (positive Bliss score) between both drugs at nanomolar concentrations (Fig. 4I; and Supplementary Fig. 3C). More importantly, AZD6738 treatment at 250 nM (highest synergistic effect) reduced more than 50% the IC50 of SN-38 in A-673 and TC-71 cells (Fig. 4J; and Supplementary Fig. 3D). Taken together, these results demonstrate that ATR inhibition enhances EwS sensitivity to SN-38.

### EWS::FLI1 promotes R-loops accumulation independently of its transcriptional regulatory activity

Based on the recently described accumulation of R-loops in EwS cells^11^, we also explored the potential contribution of EWS::FLI1-DHX9 interaction to the high levels of endogenous R-loops (independently of genotoxic agents). First, we quantified R-loops in EwS and non-EwS cells by s9.6 ICC. As previously described, we observed that EwS cells exhibited higher levels of R-loops than other sarcoma cells (Supplementary Fig. 4A). Loss of s9.6 signal after *in vitro* treatment with RNH1, but not with RNAse A, indicates the specificity of the antibody (Supplementary Fig. 4B). Importantly, the analysis of R-loop levels in a tissue microarray, including several sarcoma subtypes, corroborated that EwS samples accumulate higher levels of R-loops than other sarcomas (Fig. 5A). More importantly, downregulation of EWS::FLI1 in shA673 cells resulted in the decrease of endogenous R-loops levels, measured by s9.6 ICC (Fig. 5B). These results were confirmed by s9.6 IF and slot blot assay (Fig. 2H; Fig. 5C). *In vitro* treatment with RNH1 was used as a control of s9.6 specificity (Fig. 5C; and Supplementary Fig. 4C). Collectively, these results indicate that the excess of endogenous R-loops in EwS cells depends on EWS::FLI1.

**Figure 5.**
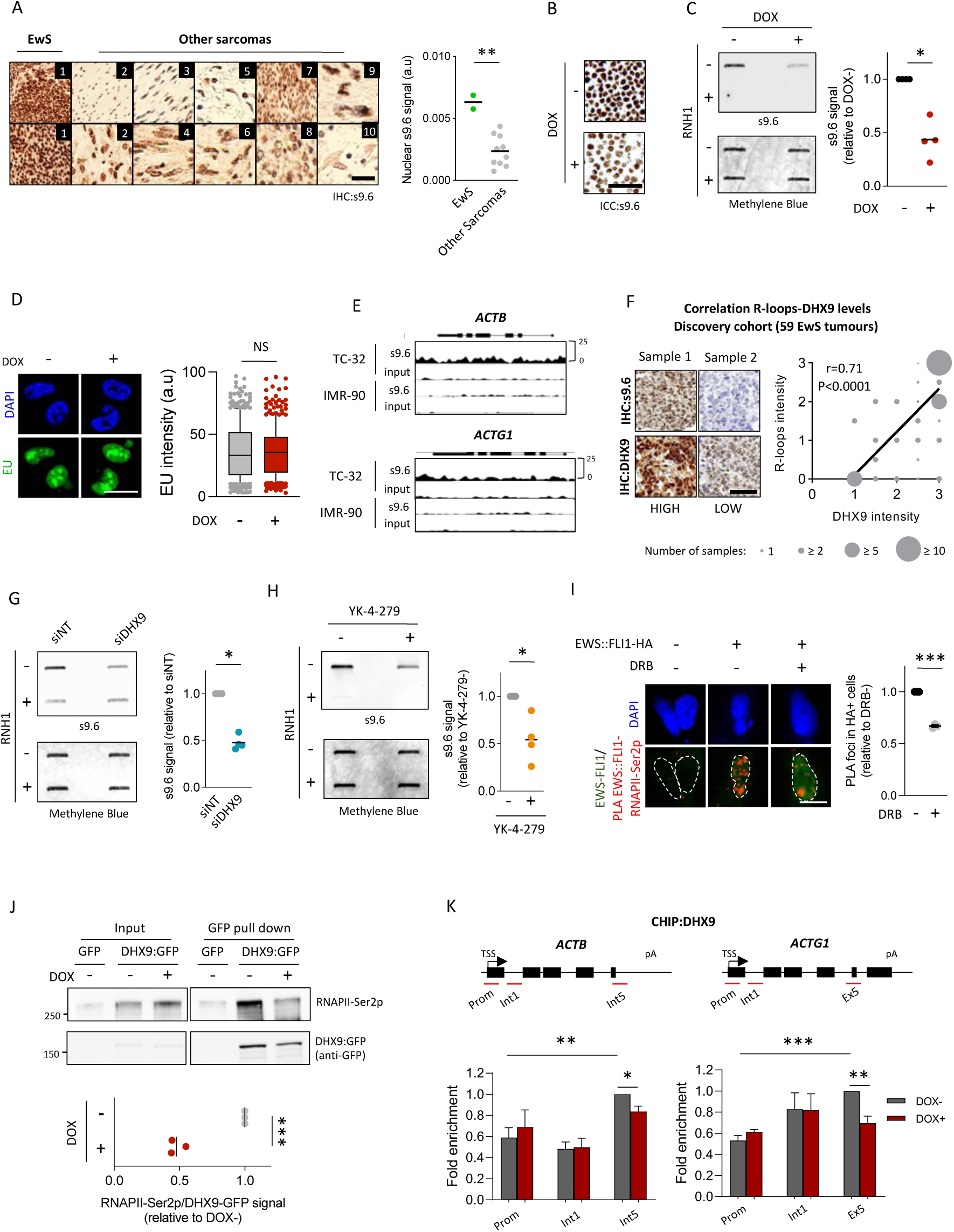
EWS::FLI1 recruits DHX9 to RNAPII transcriptional complex promoting R-loops accumulation. **(A)** Evaluation of R-loops levels in a sarcoma tissue microarray *Left*, representative images (1. EwS; 2. Osteosarcoma; 3. Synovial sarcoma; 4. GIST; 5. Differentiated Liposarcoma; 6. Undifferentiated Liposarcoma; 8. Embryonal Rhabdomyosarcoma; 9. Leiomyosarcoma; 10. Myxofibrosarcoma). Scale bar, 20 µm. *Right*, data represent mean of s9.6 nuclear signal. **(B)** Evaluation of the effect of EWS::FLI1 knockdown on R-loop levels in shA673 paraffin-embedded pellets by s9.6 ICC. Where is indicated, cells were treated with DOX for 24 h. Scale bar, 20 µm. **(C)** Similar to (B), by slot blot assay. *Left*, representative immunoblots. Loading control: methylene blue staining. *Right*, data represent mean of s9.6 signal (relative to DOX-), n=4 independent experiments. **(D)** Analysis of the effect of EWS::FLI1 knockdown in global transcription by EU incorporation. sh673 cells were incubated with DOX for 24 h. *Left*, representative images. DAPI counterstain. Scale bar, 20 µm. Right, data represent mean (±10-90 percentile) nuclear EU intensity, n=3 independent experiments. **(E)** Analysis of DRIP signal between TC-32 and IMR-90 cell lines in ACTB and ACTG1 cell lines using public dataset (GSE68845). **(F)** Analysis of the correlation between R-loops and DHX9 protein levels in a cohort of 59 EwS tumors. Correlation was determined by two-tailed Pearson correlation test. Scale bar, 50 µm. **(G)** Evaluation of the effect of DHX9 knockdown on R-loops levels by slot blot assay. *Left*, representative immunoblots. Loading control: methylene blue staining. *Right*, data represent mean of s9.6 signal (relative to siNT), n=4 independent experiments. **(H)** Evaluation of the effect of YK-4-279 treatment on R-loops levels by slot blot assay. A-673 cells were treated with 75 µM YK-4-279 (1.5 h). Other details as in (G). **(I)** Analysis of EWS::FLI1-RNAPII-Ser2p interactions by PLA using anti-HA and anti-RNAPII-Ser2p antibodies. shA673 cells overexpressing EWS::FLI1-HA were treated with 100 µM DRB (2 h). *Left*, representative images (red, PLA foci; green, EWS::FLI1-HA overexpression; blue, DAPI counterstain). Scale bar, 10 µm. *Right*, data represent mean of PLA foci number per cell (relative to DRB-), n=3 independent experiments. **(J)** Evaluation of the effect of EWS::FLI1 knockdown in DHX9-RNAPII-Ser2p interaction. shA673 cells overexpressing DHX9:GFP or control plasmids were incubated with DOX for 24 h previous to GFP pulldown. *Upper,* representative immunoblots. Molecular weight in kDa. *Bottom*, data is the mean of the RNAPII-Ser2p band signal relative to DHX9:GFP pulldown (relative to DOX-). n=3 independent experiments**. (K)** Evaluation of DHX9 recruitment to chromatin by CHIP upon EWS::FLI1 knockdown. *Upper*, diagram of *ACTB* and *ACTG1* genes; and primers used for qPCR (red lines). *Bottom,* data represent mean (±SEM) of DHX9 CHIP signal (relative to DOX-). Where is indicated, shA673 cells were treated with DOX for 24 h. Statistical significance was determined by t-test (NS, not significant; *P<0.05; **P<0.01; ***P<0.001).

**Figure 6.**
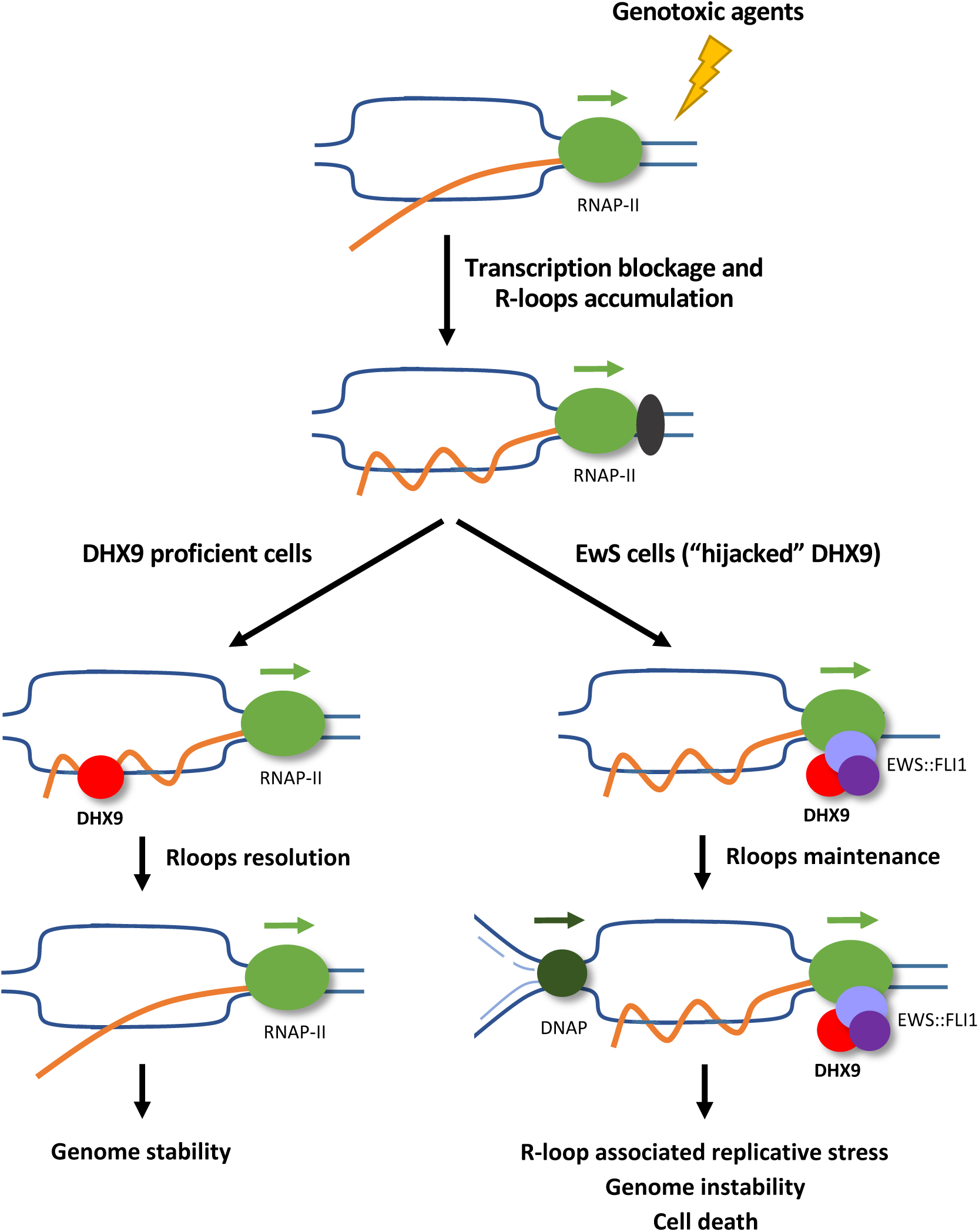
Model. Chemotherapy is a source of R-loops accumulation thought the generation of DNA adducts and the blockage of RNAPII transcription. In non-EwS cells, DHX9 resolves drug-induced R-loops preventing genome instability. In EwS cells, EWS::FLI1 “hijacks” DHX9 into the RNAPII transcriptional complex, reducing DHX9 recruitment to the R-loops and R-loops resolution. Replication machinery collapse with unresolved R-loops promoting replicative stress, genome instability and cell death.

Given the role of EWS::FLI1 as an aberrant transcription factor, we investigated whether the observed decrease in the level of R-loops upon EWS::FLI1 knockdown could result from a reduction in transcription. Interestingly, incubation of shA673 cells with DOX for 24 h did not significantly affect transcription rates, measured by EU incorporation and RNAPII-Ser2p levels (Fig. 5D; and Supplementary Fig. 4D). As a control, shA673 cells were treated with RNAPII transcription inhibitor 5,6-dichloro-1-β-D-ribofuranosylbenzimidazole (DRB), a CDK9 inhibitor that specifically blocks RNAPII elongation reducing EU incorporation and RNAPII-Ser2p levels (Supplementary Fig. 4E, 4F). To further confirm these data, we reanalyzed a published DRIP-seq dataset including EwS (TC-32, EWS-502 and CHLA-10) and non-EwS fibroblast IMR-90 cells^11^. Analysis of DRIP signal in non-EWS::FLI1 transcriptional regulated genes^39^ (*ACTB, ACTG1, CFL1* and *TPT1*) showed that EwS cells accumulated higher levels of R-loops than IMR-90 (Fig. 5E; Supplementary Fig. 5A, 5B). Importantly, no differences were observed in the expression of these genes between EwS and IMR-90 cells (Supplementary Fig. 5C). Taken together, our findings demonstrate that accumulation of R-loops in EwS is associated with EWS::FLI1 but independent of a global increase in transcription.

### EWS::FLI1 recruits DHX9 to the RNAPII transcriptional complex promoting R-loop formation

Next, we investigated whether EWS::FLI1 promotes the accumulation of endogenous R-loops in EwS cells through its interaction with DHX9. First, we evaluated the relationship between DHX9 and R-loop levels in a cohort of 59 EwS primary tumors. Results indicated a significant positive correlation (r = 0.71, P < 0.0001) (Fig. 5F). These data were confirmed in a validation cohort containing 183 EwS samples (r = 0.41, P < 0.0001) (Supplementary Fig. 6A). Interestingly, depletion of DHX9 reduced overall R-loop levels in A-673 cells (Fig. 5G). More importantly, treatment with YK-4-279 inhibitor also resulted in a significant decrease of R-loops, formally assigning to EWS::FLI1-DHX9 interaction the increased levels of R-loops in EwS cells (Fig 5H). Importantly, knockdown of DHX9 or YK-4-279 treatment did not affect cell transcription (Supplementary Fig. 6B, 6C).

While DHX9 is mainly located close to gene promoters, prolonged interaction with elongating RNAPII facilitates DHX9-dependent increase of R-loops^20^. To study possible variations in DHX9 distribution in EwS, we pulled down elongating RNAPII in A-673 cells. Notably, RNAPII-Ser2p co-precipitated DHX9 (Supplementary Fig. 6D), confirming the interaction between both proteins. In agreement, inhibition of RNAPII phosphorylation at Ser2 by DRB treatment reduced the levels of DHX9 bound to chromatin (Supplementary Fig. 6E). To evaluate whether EWS::FLI1 was implicated in the binding between DHX9 and RNAP-Ser2p, we first analyzed the interaction between EWS::FLI1 and RNAP-Ser2p by proximity ligation assay (PLA). PLA using anti-HA and anti-RNAPII-Ser2p antibodies showed an enrichment of PLA foci in A-673 cells overexpressing EWS::FLI1-HA compared to non-transfected cells (Fig. 5I). Notably, DRB treatment significantly reduced the number of foci, confirming that the interaction between both proteins is associated to transcription elongation (Fig. 5I). More importantly, knockdown of EWS::FLI1 significantly reduced DHX9-RNAPII-Ser2p physical interaction demonstrating that EWS::FLI1 increases DHX9 association to elongating RNAPII (Fig. 5J). Additionally, knockdown of EWS::FLI1 resulted in a reduction of DHX9 signal at 3’ of two constitutive genes (Fig. 5K). Altogether, these results indicate that EWS::FLI1 increases DHX9 interaction with the elongating form of RNAPII, favoring R-loop formation.

## Discussion

In this study, we conducted a comprehensive analysis of the molecular mechanisms underlying EwS sensitivity to the TOP1 poisons, which are being evaluated in clinical trials for relapsed EwS tumors^7,8^. Based on the observed hypersensitivity of EwS to TOP1 poisons, both *in vitro* and *in vivo*, we investigated the potential modulation of SN-38 activity by EWS::FLI1. Considering the intrinsic difficulty of studying EWS::FLI1 activity in EwS cell lines due to the lack of isogenic wild-type counterparts and the essential role of EWS::FLI1 in EwS cell survival, we employed a well-established doxycycline-dependent shRNA targeting EWS::FLI1^31^. This isogenic system demonstrated that EWS::FLI1 fusion is responsible for SN-38 sensitivity, highlighting the particularity of EwS biology compared to other sarcomas. Bearing in mind that genotoxic chemotherapy is the backbone of current EwS treatment regimens, these results highlighted the relevant connection between EWS::FLI1 expression and drug sensitivity for the improvement of EwS therapies.

Notably, EwS sensitivity to SN-38 was associated with a significant EWS::FLI1-dependent increase in DNA damage and genome instability. Intriguingly, SN-38-associated genome instability and sensitivity in EwS did not correlate with increased proliferation, DNA replication or transcription compared with other less sensitive sarcoma models. Furthermore, we did not observe any significant differences in the efficiency of homologous recombination repair between EwS and non-EwS sarcomas. These data suggest that SN-38 hypersensitivity is not linked to the ‘BRCAness’ phenotype previously reported in EwS cells^11^. Then we focused on R-loops due to the described accumulation of DNA:RNA hybrids upon TOP1 poisoning^33,34^. Strikingly, our data indicate that EwS cells accumulate drug-induced R-loops depending on EWS::FLI1 expression and that the enhancement of R-loop resolution by RNH1 overexpression resulted in a significant decrease in DNA damage. This result suggests that drug-induced R-loops mediate the high levels of genome instability induced by SN-38 in EwS cells.

Considering the recently described interaction between EWS::FLI1 and DHX9 and the relevance of DHX9 in the control of TOP1 poison-associated R-loops, we tested the influence of this interaction on SN-38 sensitivity. Notably, in this work, we revealed that EWS::FLI1 phenocopies DHX9 depletion. Indeed, restoration of DHX9 activity by overexpression of DHX9 or by the chemical disruption of EWS::FLI1-DHX9 interaction resulted in a reduction of TOP1-poisoning induced R-loops, DNA damage and cytotoxicity. Our results suggest that EWS::FLI1 hijacks DHX9 preventing the resolution of SN-38-induced R-loops thus promoting genome instability and cell death. Based on that, we observed a significant association between high expression levels of DHX9 and poor EwS outcome. Little is known about the mechanism that could modulate the levels and/or activity of DHX9 in EwS. Exploration of two public cohorts (n = 219 samples) indicated that the mutational frequency of DHX9 in EwS patients is about 0.5% (n = 218, cBioportal)^10,40^. High levels of DHX9, located in chr1q25, could be associated with the gain of this chromosomal arm, occurring in around 18% of EwS patients and associated with a poor patient outcome^10^.

Non-physiological R-loops pose a threat to genomic stability, especially during DNA replication. Accordingly, SN-38 exposure resulted in R-loop-associated replicative stress in EwS cells. One of the remarkable findings of this study is that replicative stress caused by SN-38 treatment was significantly dependent on EWS::FLI1 fusion, and prevented by RNAseH1 overexpression. Indeed, EwS cells showed significantly higher levels of drug-induced replicative stress compared to non-EwS ones. Importantly, we showed that the high replicative stress was promoted by EWS::FLI1-dependent defects on replication fork progression, revealing that EwS alteration of R-loops resolution contributes to the hypersensitivity to SN-38. It is important to notice that R-loop formation and TOP1 poisoning have been linked to DNA breaks induced by DNA nicks and DSBs that would favour hybridization between the RNA and the template strand^13^. However, the rapid increase of R-loops by SN-38 is in agreement with transcription-associated R-loops being transiently stabilized by TOP1 poisoning^41^. Considering the significant reduction of CHK1 phosphorylation induced by SN-38 upon EWS::FLI1 knockdown, we suggest the latter as the most likely explanation. Interestingly, inhibition of replication fork stress signalling by ATR inhibitor cerelasertib exacerbated SN-38 sensitivity. Altogether these results open the possibility of combined therapy using genotoxic chemotherapy and ATR inhibitors which have been previously tested in single therapy^42^.

Considering the dual role of DHX9 in the metabolism of R-loops we asked about the relationship between EWS::FLI1-DHX9 interaction and the high levels of endogenous R-loops in EwS compared to other sarcomas (Fig. 5A). Indeed, we demonstrated that these high endogenous R-loop levels also depended on EWS::FLI1-DHX9 interaction and that EWS::FLI1 phenocopies DHX9 depletion. EwS cells showed fusion-dependent interaction of DHX9 and the elongating form of RNAPII, supporting a model in which EWS::FLI1 promotes DHX9 association with elongating RNAPII, similar to the pattern promoted by defects on splicing^20^. Importantly, we corroborated this connection by observing that high levels of DHX9 significantly correlated with high levels of R-loops in EwS patients. Further research is required to validate this uncovered mechanism and to study the possible role of endogenous R-loops in EwS biology.

Overall, our results demonstrate that EWS::FLI1 modulates the activity of DHX9 in the metabolism of R-loops, leading to an increase in endogenous R-loops and preventing the resolution of those induced by SN-38. The consequence of this is an increased sensitivity of EwS cells to TOP1 inhibitors. These findings expand our knowledge of the mechanisms conferring sensitivity of EwS cells to genotoxic agents, which could contribute to the future development of novel combinatorial therapies.

## Methods

### Cell lines and culture conditions

Human EwS: A4573, A-673, TC-71 and EW-7; human breast cancer: MBA-MB-436 and HCC-1937, human fibroblast MRC-5 and sarcoma derived U2OS, HT-1080, SK-UT-1 and 93T449 cells lines were purchased from ATCC. A-673/TR/shEF^30^ cell line was gently provided by J. Alonso (Madrid, Spain). All cell lines were grown at 37 °C, 5% CO_2_, routinely tested for mycoplasma contamination using the MycoAlert Detection Kit (Lonza Group Ltd, LT07-318) and authenticated by STR analysis (CLS, Cell Lines Service GmbH). Cell lines characteristics, origin and culture conditions are depicted in <u>Supplementary Table 1</u>. A-673/TR/shEF cells were incubated with 1 µg/ml doxycycline (DOX) (Sigma, D9891) for indicated times. Drugs used in this study are listed in <u>Supplementary Table 1</u>.

### Patient-Derived Xenograft (PDX) models

EwS (IEC-73, HSJD-006, HSJD-002) and non-EwS (osteosarcoma, IEC-036; undifferentiated pleomorphic sarcoma, IEC-056) PDXs were subcutaneously implanted in nu/nu athymic nude mice (Envigo) in both flanks. When tumor volume was around 100 mm^3^, mice were randomized in two groups and 10 mg/kg irinotecan (Accord Healthcare, 713386.5) was injected intraperitoneally 5 times per week during 2 weeks. Control group was treated with physiological saline solution. Tumor size and mice weight were measured every two days. Tumor volume was calculated as *V*= (*a* x *b*^2^ x ν)/6, where *b* is the smallest measure and *a* the largest measured. Part of animals were sacrificed at day 7 of treatment and tumors were embedded in paraffin for the evaluation of tumoral necrosis. Hematoxylin/Eosin staining was performed according to conventional protocols and evaluated by an experienced pathologist. When tumor volume reached the predefined humane endpoint of around 1,200 mm3 (= event) animals were sacrificed by cervical dislocation. Animal experiments were approved by the Consejeria de Agricultura, Pesca, Agua y Desarrollo Rural; Junta de Andalucia (08-05-2018-082) and performed according to protocols and conditions reflected in the European guidelines (EU Directive 2010/63/EU).

### Immunocytochemistry (ICC) and Immunohistochemistry (IHC)

For ICC, cell lines were collected and pellets were fixed in 25% formalin before inclusion in paraffin. ICC and IHC were performed at HUVR Biobank facility according to conventional protocols using antibodies listed in <u>Supplementary Table 1</u>. Briefly, 4 µm tissue sections from paraffin blocks were dewaxed in xylene and rehydrated in a series of graded alcohols. Antigen retrieval was performed with a PT link instrument (Agilent) using EDTA buffer (pH 8.0). After that, sections were incubated in H_2_O_2_ for 30 min and blocked with 1% blocking solution (Roche) in PBS-0.05% Tween-20. After primary antibodies, sections were incubated with peroxidase-labeled secondary antibodies and 3,3-diaminobenzimide, according to manufacturer protocol (Leica Biosystems). Slides were counterstained with Hematoxylin and mounted using DPX (BDH Laboratories). Samples were evaluated by an experience pathologist and scored according to the staining intensity in: 0 (negative), 1 (low), 2 (moderate) and 3 (high).

### siRNA and plasmid transfection

Transient cell transfections with siRNA and plasmids were performed using jetPRIME (Polyplus, 101000046) according to the manufacturer protocol. siRNAs and plasmids are listed in <u>Supplementary Table 1</u>.

### Western blotting

Western blotting was performed as previously described by our group^43^. Briefly, cell lines were lysed using RIPA buffer (10 mM Tris-HCl pH 8, 1 mM EDTA, 0.5 mM EGTA, 1% Triton X-100, 0.1% Sodium Deoxycholate, 0.1 % SDS, 140 mM NaCl), supplemented with protease (Roche, 11697498001) and phosphatase inhibitors (1 mM Na_3_O_4_V (Sigma-Aldrich, 450243)) and 1 mM NaF (Sigma-Aldrich, S1504)). Protein concentration was determined using Pierce^TM^ BCA Protein Assay Kit (Thermofisher, 23225). Primary antibodies (listed in <u>Supplementary Table 1</u>) were blocked in Tris buffered saline buffer 0.1% Tween-20; 5% BSA. HPRT-conjugated anti-mouse (Biorad, 1706516) and anti-rabbit (Biorad, 1706515) secondary antibodies were incubated for 1 h at room temperature (1:5000). Images were capture using a Chemidoc Imaging System (Bio-Rad) and band density was determined using the Image Lab Software (Bio-Rad).

### Subcellular fractionation

For the analysis of chromatin-binding proteins, cells were grown in 100 mm cell culture dishes up to 80-90% of confluency. After treatment with indicated drugs, cells were washed twice with ice-cold PBS and collected through scrapping. 10% of cells (WCE) were lysed in 2x LB (125 mM Tris-HCl pH 6.8, 4% SDS, 0.02% bromophenol blue, 20% glycerol, 200 mM DTT) and maintained on ice. The rest of cells were lysed with Lysis Buffer 1 (250 nM sucrose, 50 mM Tris-HCl, 5 mM MgCl_2_, 0.25% NP40) on ice for 10 min. After centrifugation, nuclei were washed with Buffer 2 (1 M sucrose, 50 mM Tris-HCl, 5 mM MgCl_2_) and lysed using Nuclear Buffer Lysis 3 (20 mM Tris-HCl, 0.4 M NaCl, 15% glycerol, 1.5% Triton X-100, 100 mM CaCl_2_). After centrifugation, chromatin fraction was resuspended in 2x LB and boiled (with WCE) twice at 99 °C for 5 min. Protein extracts were finally sonicated in a Bioruptor (Diagenode) for 3 cycles at high intensity. Western blotting was performed as previously described.

### Coimmunoprecipitation (co-IP)

For RNAPII-Ser2p immunoprecipitation, a subcellular fractionation was performed, as previously described. Nuclei lysates (in Nuclear Buffer Lysis 3, supplemented with protease (Roche, 11697498001) and phosphatase inhibitors (1 mM Na_3_O_4_V (Sigma-Aldrich, 450243)) and 1 mM NaF (Sigma-Aldrich, S1504)), were sonicated and incubated overnight under rotation at 4 °C with 5 ug/ml of anti-RNAPII-Ser2p (Abcam, ab193468) or anti-IgG (Cell Signaling, 3900). Next, extracts were incubated with Dynabeads^TM^ Protein G (Invitrogen^TM^, 10004D) for 2 h under rotation at 4 °C. After washes, beads were resuspended in 2x LB and boiled at 99 °C for 5 min. Western blotting was performed as previously described. For DHX9:GFP immunoprecipitation, cells were collected upon 48 h DHX9:GFP or control plasmid transfection. Next, cells were lysed in Lysis Buffer (25 mM HEPES pH 8, 150 mM NaCl, 10% Glycerol, 0.5% Triton X-100) and sonicated for 3 cycles at low intensity. Protein extracts were incubated with ChromoTek GFP-Trap Agarose beads (ProteinTech, GTA-20) under rotation for 2 h at 4 °C. After washes, beads were resuspended in 2x LB and boiled at 99 °C for 5 min. Western blotting was performed as described above.

### Immunofluorescence

Cells were cultured in fibronectin-coated coverslips for 24 h before treatment or transfection. After that, slides were washed twice with PBS and fixed with 4% PFA (Thermo Scientific, 28908) for 10 min at room temperature. Next, cells were permeabilized with PBS-0.2% Triton X-100 for 10 min, blocked with PBS-5% BSA for 1 h and incubated with primary antibodies for 1-3 h in PBS-1% BSA. Primary antibodies used in this study are listed in <u>Supplementary Table 1</u>. Slides were washed twice with PBS-1% BSA for 10 min and incubated with corresponding Cy2/Cy3-conjugated secondary antibodies (Jackson Laboratories) for 1 h. After washes, cells were counterstained with DAPI (Sigma-Aldrich, 09542) for 5 min and mounted using Dako Fluorescent Mounting Medium (DAKO, S3023). Images were taken using Olympus BX61 Fluorescence Motorized Microscope (Olympus) and analyzed using ImageJ software.

### Cell cycle and apoptosis analysis by flow cytometry

For cell cycle analysis, cells were washed twice with PBS and fixed with 70% ethanol overnight at 4 °C. Then, cells were washed twice with PBS and incubated with propidium iodide (PI) solution (100 µg/ml PI and 100 µg/ml RNAse A) for 30 min. Finally, cells were washed with PBS before FACS acquisition. For the analysis of apoptosis, cells and supernatant were collected and washed twice with PBS. After that, cells were incubated with AnnexinV and PI (Immunostep; FITC-conjugated, ANXVKF7-100T or DY634-conjugated, ANXVKDY-100T) according to the manufacturer instructions for 15 min. 10,000-20,000 cells were counted for each sample using a BD FACSCanto II flow cytometer (Becton Dickinson). Data were analyzed using FlowJo v.10.2 software and total AnnexinV positive cells (early + late apoptosis) were shown.

### Metaphase spreads and sister chromatid exchanges (SCEs) assay

SCEs assay was performed as previously described by our group^44^. Cells were incubated with 10 μM BrdU (Sigma-Aldrich, B5002) for 40 h. Then, cells were treated with 2.5 µM etoposide (Sigma-Aldrich, E1396) for 30 min, and washed twice with PBS. After treatment, cells were cultured in drug-BrdU-free, 0.2 μg/ml demecolcide-containing media (Sigma-Aldrich, D1925) for 8-14 h. Cells were collected and incubated in 0.03 M trisodium citrate solution for 30 min at 37 °C. After that, cells were fixed in 3:1 methanol:acetic acid. Fixed cells were dropped onto acetic acid-humidified slides and incubated in 10 μg/ml bisbenzimide solution (Hoechst 33258, Sigma-Aldrich, 14530) for 20 min. Next, slides were washed in 20x SSC buffer and irradiated in a 365 nm UV-lamp for 45 min. Finally, slides were incubated in 20x SSC at 65 °C and stained for 20 min with 5% Giemsa solution (Sigma-Aldrich, GS500).

### RNA extraction, reverse transcription and quantitative real-time PCR

Total RNA was extracted from cultured cells using the miRNeasy Kit (Qiagen, 217084) according to the manufacturer’s instructions. DNA was removed using RQ1 DNAse (Promega, M6101). RNA quantity was measured using a Nanodrop ND-2000 Spectrophotometer (Thermo Scientific). 500-1000 ng of RNA were retrotranscribed using MultiScribe Reverse Transcriptase Kit (Invitrogen, 4311235) according to manufacturer protocol. qPCR was performed on a 7900HT Fast Real-Time PCR system (Applied Biosystems) using TaqMan probes and TaqMan Universal PCR Master Mix (Applied Biosystems, 4326708). TaqMan probes are listed in <u>Supplementary Table 1</u>.

### EdU and EU incorporation assay

Cells grown on coverslips were incubated with 0.5 mM EU for 45 min (BaseClick, BCN-003) or 10 µM EdU for 15 min (BaseClick, BCN-001). Next, coverslips were washed twice with PBS and fixed with 4% PFA for 10 min at room temperature. Cells were permeabilized with PBS-0.2% Triton X-100 for 10 min and blocked with PBS-5% BSA for 1 h. After washout, cells were incubated with the Reaction cocktail solution (100 mM Tris-HCl pH 8.5, 1 mM CuSO_4_, 1 µM Fluorescein Azide 6-FAM (BaseClick, BCFA-001), 100 mM ascorbic acid) for 30 min at room temperature, under agitation and in the dark. After incubation, coverslips were washed twice with PBS-1% BSA and 4 times with PBS-0.1% Tween 20 for 10 min. Finally, cells were counterstained with DAPI for 5 min and mounted using Dako Fluorescent Mounting Medium. Images were taken using Olympus BX61 Fluorescence Motorized Microscope (Olympus) and analyzed using ImageJ software.

### Proximity ligation assay (PLA)

PLA was performed according to manufacturer instructions (Sigma-Aldrich, Duolink In Situ Detection Reagents Red, DUO92008). Briefly, cells grown on coverslips were fixed with 4% PFA for 10 min at room temperature and permeabilized with PBS-0.2% Triton X-100 for 10 min. Primary antibodies (listed in <u>Supplementary Table 1</u>) were incubated for 1.5 h at room temperature. Finally, cells were counterstained with DAPI for 5 min and mounted using Dako Fluorescent Mounting Medium. For EWS::FLI1-HA/RNAPII-Ser2p PLA, coverslips were additionally incubated with Cy2-conjugated anti-mouse secondary antibody for 1 h, before DAPI counterstained, for the detection of EWS::FLI1 overexpressing cells.

### Slot blot

Slot blot was carried out as previously described^45^ with modifications. Cells were lysed overnight at 37 °C by using SDS/Proteinase K standard method. Next, nucleic acids were extracted by phenol/chloroform and EtOH/sodium acetate purification. Pellets were resuspended in mQH_2_O and sonicated in a Bioruptor (Diagenode) for 3 cycles at high intensity. 1 µg of DNA was treated with 0.1 ng of RNAse A (Invitrogen, AM2271) at 37 °C for 2 h. For RNAse H1 controls, 1 ug of DNA was incubated with 6U RNAse H (NEB, M0297) at 37 °C for 2 h. Then, samples were loaded into a H_2_O pre-humidified Hybond-N+ Nylon membrane (Amersham, RPN303B) using a slot blot apparatus (Hoefer, PR648). Membranes were UV-crosslinked (0.12 J/m^2^), blocked with TBST-5% non-fat milk at room temperature for 1 h and incubated with 1:1000 s9.6 (Kerafast, ENH001) overnight at 4 °C. After washes, membranes were incubated with 1:5000 HPRT-conjugated anti-mouse secondary antibody (Biorad, 1706516) for 1 h at room temperature. After acquisition of images using a Chemidoc Imaging System (Bio-Rad), membranes were incubated with 0.02% Methylene blue in 0.3 M sodium acetate pH 5.2 for 10 min to stain total DNA.

### DNA fibers

DNA fiber assay was performed as previously described^46^. Briefly, cells were labeled with 25 µM 5-Chloro-2’-deoxyuridine (CldU, Sigma-Aldrich, CAS 50-90-8) for 30 min. After washes, cells were incubated with 50 µM 5-Iodo-2’-deoxyuridine (IdU, Sigma-Aldrich, CAS 54-42-2) and 5 µM SN-38 for 30 min. Then, cells were collected and dropped on frosted slides. Immediately, cells were lysed with Spreading Buffer (200 mM Tris-HCl pH 7.5, 50 mM EDTA pH 8, 0.5% SDS) and fixed with methanol:acetic acid solution 3:1. Next, DNA was denatured with 2.5 M HCl for 1 h and incubated with 1:1000 rat anti-BrdU (Abcam, ab6326) for 1 h at 4 °C, with 1:500 donkey anti-rat Alexa Fluor 488 secondary antibody (Thermofisher, A21208) for 1.5 h at 4 °C, with 1:500 mouse anti-BrdU (BD biosciences, 347580) overnight at 4 °C and, finally, with 1:250 goat anti-mouse Alexa Fluor 546 secondary antibody (Thermofisher, A11003) for 2 h at 4 °C. Montage was performed using Fluoromount-G Mounting Medium (eBioscience, 00-4958-02). Images were taken using Olympus BX61 Fluorescence Motorized Microscope (Olympus) and analyzed using ImageJ software.

### Chromatin immunoprecipitation (CHIP-qPCR)

CHIP experiments were performed using the SimpleChIP Enzymatic Chromatin IP Kit (Cell Signaling, 9003) according to manufacturer instructions. Cells were cultured in 150 mm cell culture dishes up to 80-90% of confluency. For chromatin fragmentation, extracts were incubated with 2 µl Micrococcal Nuclease (Cell Signaling, 10011) and sonicated in a Bioruptor (Diagenode) for 2 cycles at high intensity. Extracts were incubated with 5 ug of anti-DHX9 (Abcam, ab26271) or anti-IgG (Cell Signaling, 2729) overnight at 4 °C. qPCR was performed on a 7900HT Fast Real-Time PCR System (Applied Biosystems) using primers and iTaq Universal SYBR Green Supermix (Biorad, 1725270). Primers^18^ are listed in <u>Supplementary Table 1</u>.

### Survival analysis

For the correlation between DHX9 mRNA levels and EwS patient outcome, survival analysis was performed as previously described^47^ in a cohort of 166 primary EwS tumors with available clinical annotations. Patients were grouped according to DHX9 mRNA levels in: high (first quartile) and moderate-low (second to last quartile). For the correlation between R-loops levels and EwS patient outcome, s9.6 IHC was performed in a cohort including 213 EwS tumors with available clinical annotations. Patients were stratified according to R-loops levels in: high (intensity > 2) and moderate-low (intensity ≤ 2). P value was determined by Mantel-Cox test using GraphPad PRISM version 9 (GraphPad Software Inc., CA, USA).

### MTT assay and drug combinations

Cells were seeded in a 96 well cell culture plates at a density of 1.5 – 3 x 10^3^ cells per well. To analyze the sensitivity to SN-38 (Biosynth, FE29579), cells were treated with indicated concentrations of SN-38 or DMSO for 72 h before MTT assay (Roche, 11465007001), according to the manufacturer instructions. For the combination of SN-38 with DOX or YK-4-279 inhibitor (MedChemExpress, HY-14507), cells were pre-treated with 1 µg/ml of DOX or 75 µM of YK-4-279 for 24 h or 1 h, respectively. Then, cells were treated with indicated concentrations of SN-38 for 3 h, washed and maintained in drug-free medium for 48 h before MTT assay. Absorbance was measured in a microplate reader (TECAN) at 565 nm. Survival curves, IC50 and AUC was determined using GraphPad PRISM version 9 (GraphPad Software Inc., CA, USA).

For SN-38 and ATRi combination, cells were treated with AZD6738 ATRi Ceralasertib (MedChemExpress, HY-19323A) for 24 h at indicated concentrations before treatment with SN-38. Cells were maintained in drug-containing medium during 72 h previous to MTT assay. Absorbance was measured in a microplate reader (TECAN) at 565 nm. Synergy between SN-38 and ATRi was evaluated using SynergyFinder 3.0^48^.

### Analysis of microarray, DRIP-seq and RNA-seq public data

Microarray data (GSE176190)^39^ were analyzed using GEO2R. DRIP-seq data (GSE68847)^11^ were analyzed using Galaxy web-based platform. Briefly, reads were mapped against hg19 human reference genome using Bowtie2 and peaks were determined using MACS2. Data were visualized using UCSC genome browser (University of California). RNA-seq data (GSE68836)^11^ were analyzed using Galaxy web-based platform. After quality control, reads were mapped against hg19 human reference genome using Bowtie2 tool. Results were visualized using Integrative genomics viewer (IGV).

### Statistical analysis

Statistical analysis was performed using GraphPad PRISM version 9 (GraphPad Software Inc., CA, USA). Comparison tests are indicated in figure legends. Statistical significance is indicated by: NS, not significant; *P<0.05; **P<0.01; ***P<0.001).

## Acknowledgements

We thank Belén Gómez-González for critical reading of the manuscript. We also thank the support of Xarxa de Bancs de Tumors de Catalunya (XBTC) sponsored by Pla Director d’Oncologia de Catalunya.

## Author contribution

J.O-P. performed most of the experiments unless indicated, analyzed data and generated the figures. E.G-C. performed Co-IP and DNA fibers assays. D. D-B. helped in Co-IP and ChIP assays. L.L-S. helped with *in vitro* experiments. C.J-P. supported animal experiments. A.T.M-A. designed experiments and helped to write the manuscript. A.C.E. and S.O. provided microarray data. J.A. provided cellular models. AM. C. provided EwS PDX models. I.M. and A.L-B. provided tissue microarrays. T.G.P.G. provided EwS patients microarray expression data and clinical information, provided laboratory infrastructure and financial support for J.O-P. international internship and biological guidance. E. dA., F. G-H and J.O-P designed experiments and wrote the manuscript. E. dA. and F.G-H coordinated and supervised the study. All authors reviewed the manuscript.

