## Supplementary figures for "EWS::FLI1-DHX9 interaction promotes Ewing sarcoma sensitivity to DNA topoisomerase 1 poisons by altering R-loop metabolism"

**Supplementary Figure 1**. **(A)** TOP1 poisons/inhibitors effect between EwS (EWSR1::FLI1) and non-EwS cell lines from Genomics of Drug Sensitivity in Cancer website. **(B)** Evaluation of response to treatment (CR, complete response; PR partial response; SD, stable disease; PD, progressive disease) of EwS and non-EwS PDXs 21 days after the beginning of irinotecan treatment. **(C)** Kaplan-Meier curves comparing survival of PDXs models treated with irinotecan. **(D)** Evaluation of proliferation of EwS, non-EwS and MRC-5 cell lines by Ki-67 ICC in paraffin-embedded pellets. Scale bar, 50 µm. **(E)** Determination of transcriptional rates between EwS and non-EwS cell lines by EU incorporation. DAPI counterstain. Scale bar, 50 µm. **(F)** Data represent mean of SCEs induction upon 2.5 µM etoposide treatment (30 min), n=3 independent experiments. Statistical significance was determined by t-test (NS, not significant; *P<0.05; **P<0.01; ***P<0.001).

**Supplementary Figure 2**. **(A)** Evaluation by qPCR of *EWS::FLI1* and *LOX* mRNA levels in shA673 cells after incubation with DOX for the indicated hours. Data represent mean (±SEM), n=3 independent experiments. **(B)** Analysis of cell cycle of shA673 by PI FACS. Data represent the mean (±SEM) of cell cycle populations after incubation with DOX for indicated hours. **(C)** Determination of replication activity in shA673 cells after DOX incubation by EdU incorporation. *Left*, representative images. DAPI counterstain. Scale bar, 50 µm. *Right*, data represent mean of nuclear EdU intensity in EdU+ cells, n=2 independent experiments. **(D)** Evaluation of RHN1 levels in A-673 cells after RNH1:GFP or control plasmid transfection by IF. Representative images. DAPI counterstain. Scale bar, 20 µm. **(E)** Analysis of DHX9 levels in A-673 after 48 h of siRNA silencing by WB. Loading control: α-tubulin. Molecular weight in kDa**. (F)** Evaluation of DHX9 levels in A-673 and TC-71 cells after DHX9:GFP or control plasmid transfection by IF. Representative images. DAPI counterstain. Scale bar, 10 µm. Statistical significance was determined by t-test (NS, not significant; *P<0.05; **P<0.01; ***P<0.001). **(G)** Kaplan-Meier showing event-free survival of 85 EwS samples (GSE63157) stratified by DHX9 mRNA levels. Statistical significance was determined by Mantel-Cox test.

**Supplementary Figure 3**. **(A)** Analysis of the effect of EWS::FLI1 knockdown on SN-38 induced pRPA32 (s4+s8) by IF. *Left*, representative images. DAPI counterstain. Scale bar, 20 μm. *Right,* data represent mean (±SEM) of pRPA32 (s4+s8) intensity (relative to SN-38), n=3 independent experiments. **(B)** Representative immunoblots of the effect of DHX9 overexpression on SN-38-induced replicative stress in TC-71. *Bottom*, quantification of CHK1 phosphorylation (pCHK1/CHK1 band signal). Data are the mean of 2 independent experiments (relative to SN-38). Loading control: GAPDH. Molecular weight in kDa. **(C)** Dose-response matrix of the combination between AZD6738 and SN-38 in A-673 and TC-71 cell lines. **(D)** IC50 data from MTT assay. Cells were incubated with AZD6738 (250 nM, 24 h) previous to the treatment with SN-38 (72 h).

**Supplementary Fig 4. (A)** Evaluation of R-loop levels in EwS and non-EwS paraffin-embedded cellular pellets by s9.6 ICC. *Left*, representative images. Scale bar, 50 μm. *Right*, data represent mean of nuclear s9.6 signal. **(B)** Analysis of R-loops levels in shA673 paraffin-embedded cellular pellets after in vitro treatment with RNAse A or RNH1 by s9.6 ICC. Scale bar, 50 μm. **(C)** S9.6 slot blot. shA673 extracts were treated *in vitro* with RNH1 and loaded into Nylon membrane at different concentrations. Membranes were incubated with s9.6 antibodies and stained with Methylene blue. **(D)** Analysis of the effect of EWS::FLI1 knockdown in cell transcription by RNAPII-Ser2p ICC. shA673 cells were incubated with DOX for 24 h, previous to the generation of paraffin-embedded pellets. Scale bar, 50 μm. **(E)** Evaluation of the effect of DRB treatment in cell transcription by EU incorporation. shA673 cells were treated with 100 μM DRB for 2 h. **(F)** Similar to (E), by RNAPII-Ser2p ICC.

**Supplementary Fig 5. (A)** Analysis of the expression levels of *ACTB, ACTG1*, *CFL1* and *TPT1* genes in 15 EwS cell lines upon EWS::FLI1 knockdown using public dataset (GSE176190). Data represent the mean of mRNA levels in EWS::FLI1 low condition, relative to EWS::FLI1 high. **(B)** Evaluation of DRIP signal in *ACTB*, *ACTG1, CFL1* and *TPT1* genes between EwS (TC-32, EWS-502 and CHLA-10) and non-EwS IMR-90 cell lines using public dataset (GSE68845).

**(C)** Evaluation of the expression of *ACTB, ACTG1, CFL1* and *TPT1* genes between EwS (TC-32, EWS-502 and CHLA-10) and non-EwS IMR-90 cell lines using public dataset (GSE68836).

**Supplementary Fig 6. (A)** Analysis of the correlation between R-loops and DHX9 protein levels in a cohort of 183 EwS tumors. Correlation was determined by two-tailed Pearson correlation test. **(B)** Evaluation of the effect of DHX9 knockdown on transcriptional activity by EU incorporation. Left, representative images. DAPI counterstain. Scale bar, 20 µm. *Right,* data represent the mean of EU intensity (relative to siNT), n=2 independent experiments. **(C)** Similar to (B). A-673 cells were treated with 75 µM YK-4-279 (1.5 h). **(D)** Analysis of the interaction between DHX9 and RNAPII-Ser2p by RNAPII-Ser2p pulldown. Molecular weight in kDa. Asterisk indicates non-specific bands. **(E)** Evaluation of the effect of DRB treatment in the binding of DHX9 to chromatin. Left, representative plot. Molecular weight in kDa. Right, data represent the mean (±SEM) of DHX9 band intensity in the chromatin fraction (relative to DRB-), n=3 independent experiments. Statistical significance was determined by t-test (**P<0.01).

#### Supplementary Figure 1.

A

**Drug activity**
**(EWSR1::FLI1+ vs. EWSR1::FLI1- cells)**
Genomics of Drug Sensitivity in Cancer (GDSC2)

| Drug | Target | Drug effect | p-value |
|---|---|---|---|
| Irinotecan | TOP1 | -1.57 | 4.48e$^{-8}$ |
| Topotecan | TOP1 | -1.2 | 5.03e$^{-5}$ |
| Camptothecin | TOP1 | -1.11 | 6.31e$^{-5}$ |
| SN-38 | TOP1 | -0.93 | 0.006 |
| LMP744 | TOP1 | -1.11 | 0.0002 |

C

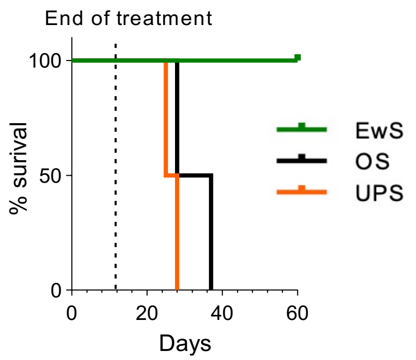


B

| Model | Disease | Nº of tumors | CR (%) | PR (%) | SD (%) | PD (%) |
|---|---|---|---|---|---|---|
| IEC-073 | EwS | 6 | 100 (6/6) | 0 | 0 | 0 |
| HSJD-ES-002 | EwS | 8 | 87,5 (7/8) | 12,5 (1/8) | 0 | 0 |
| HSJD-ES-006 | EwS | 6 | 100 (6/6) | 0 | 0 | 0 |
| IEC-036 | Osteosarcoma | 4 | 0 | 25% (1/4) | 75% (3/4) | 0 |
| IEC-056 | UPS | 4 | 0 | 0 | 100% (4/4) | 0 |

D

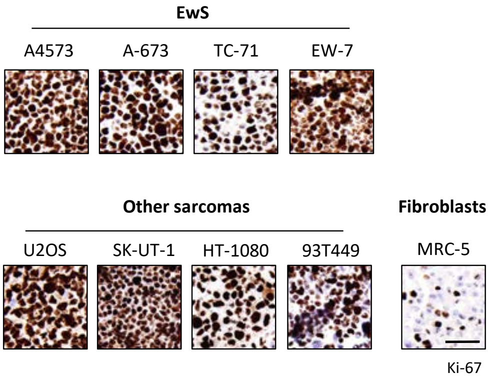


E

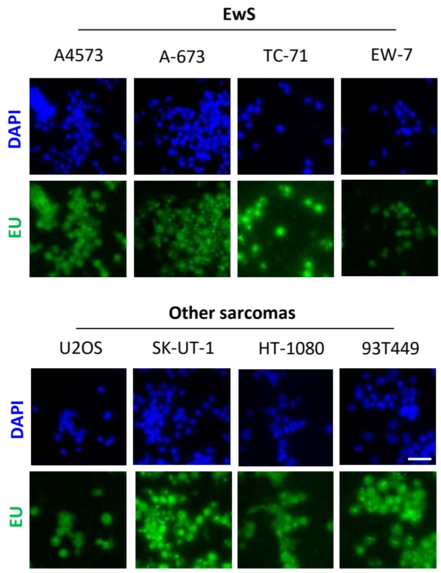


F

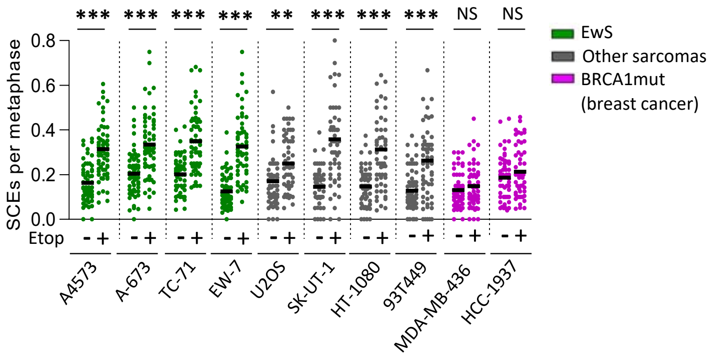

#### Supplementary Figure 2.

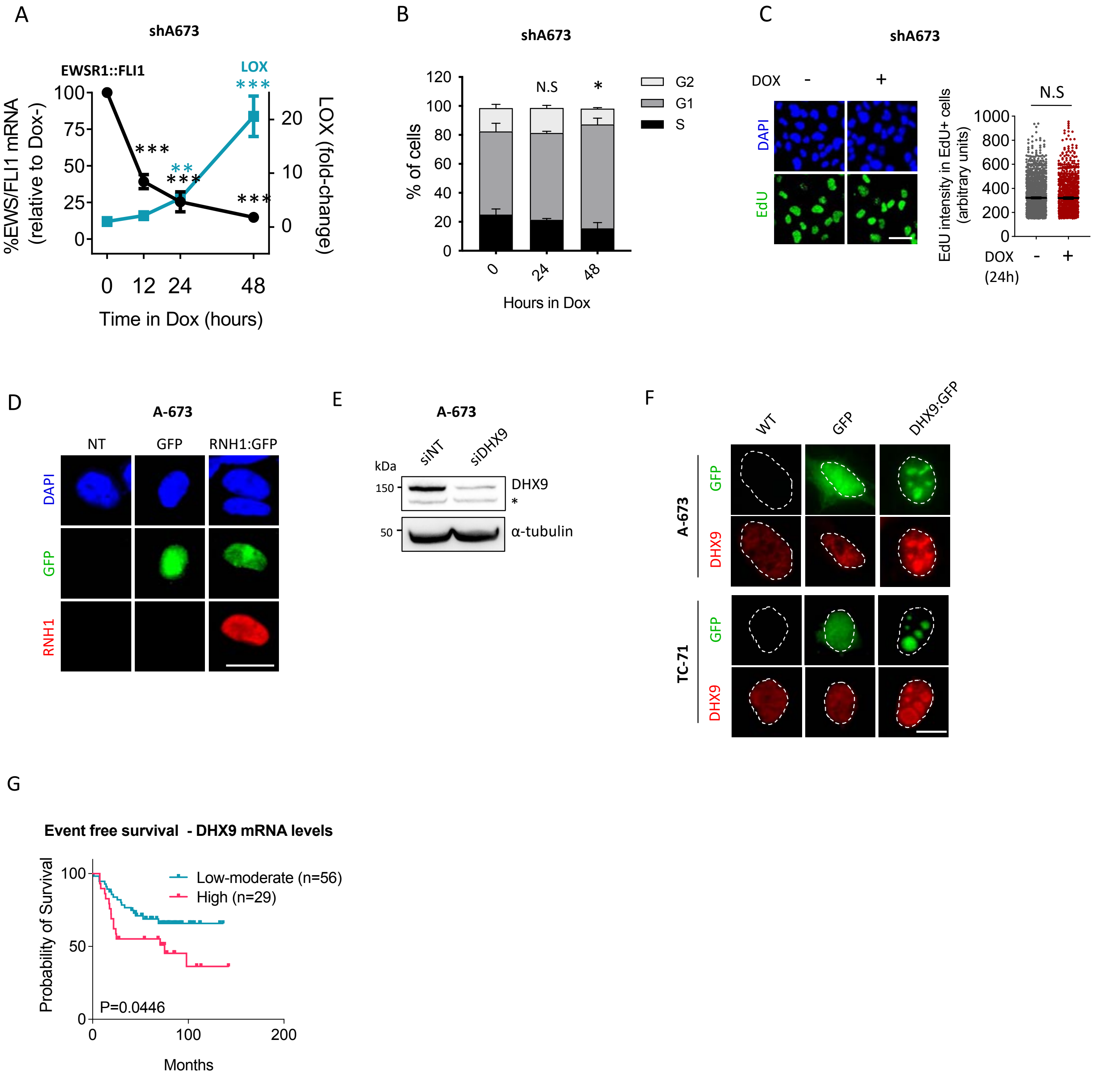

#### Supplementary Figure 3.

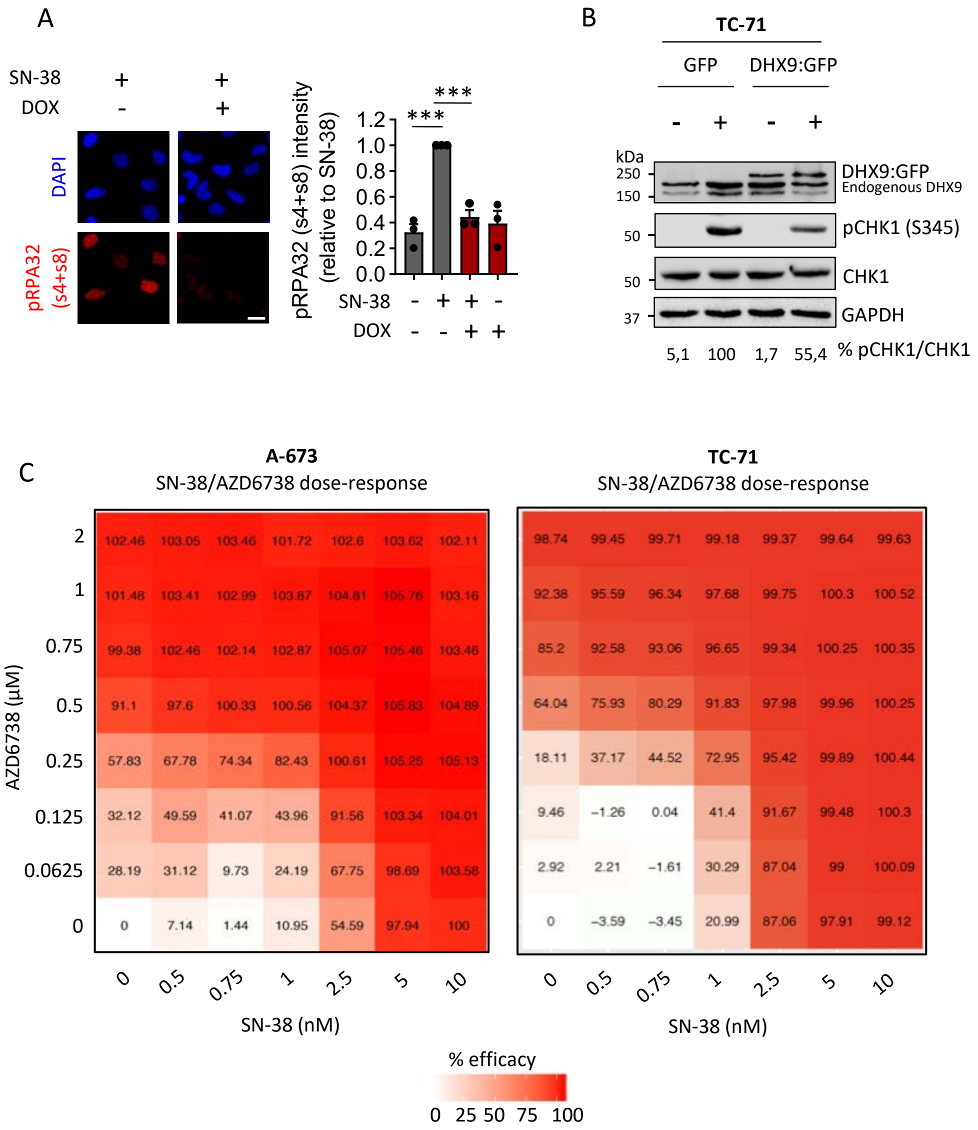


D

| | SN-38 IC50 (nM) | | |
|---|---|---|---|
| | AZD6738 - | AZD6738 + | SN-38 IC50 (AZD6728-/+) |
| **A-673** | 2.43 | 0.91 | **2.67x** |
| **TC-71** | 1.3 | 0.67 | **1.34x** |

#### Supplementary Figure 4.

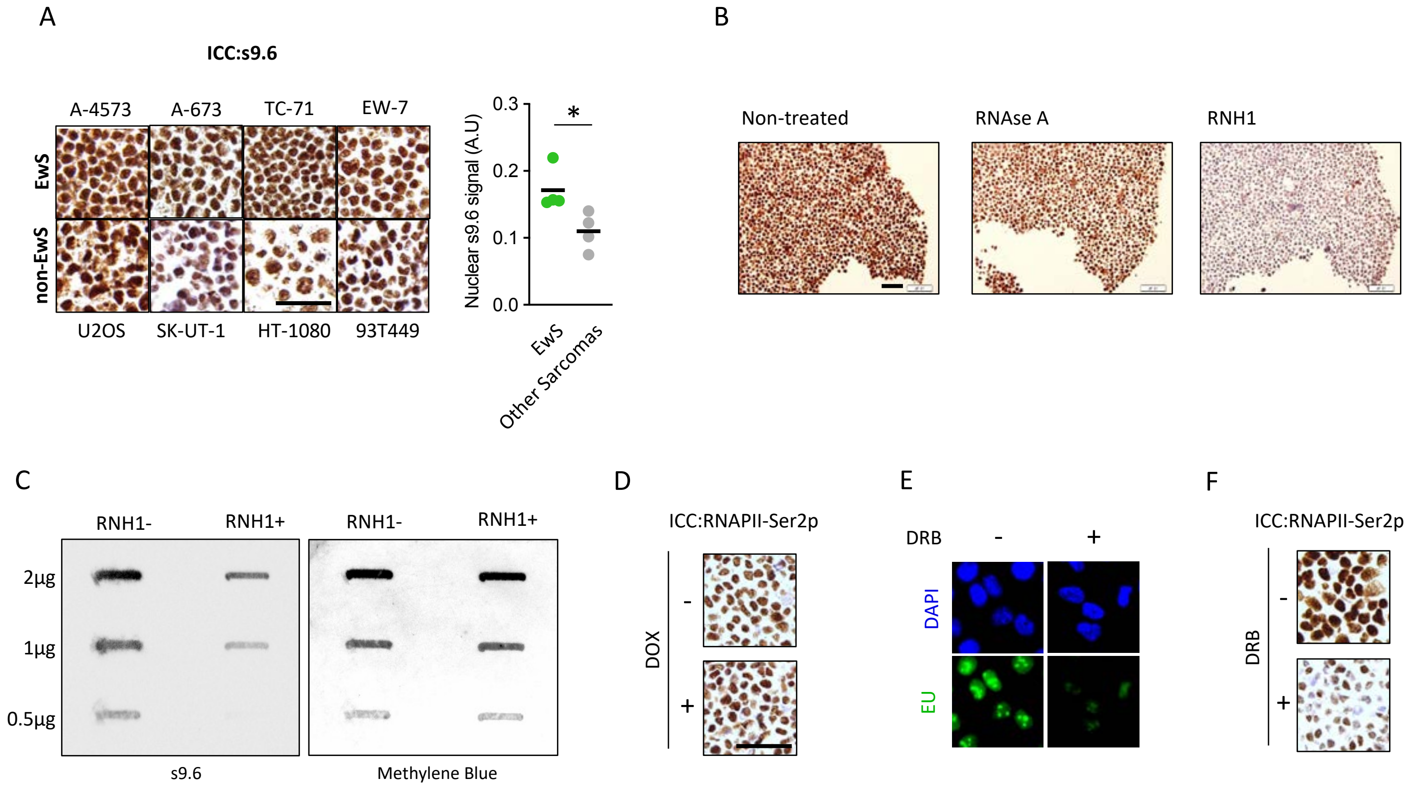

### Supplementary Figure 5.

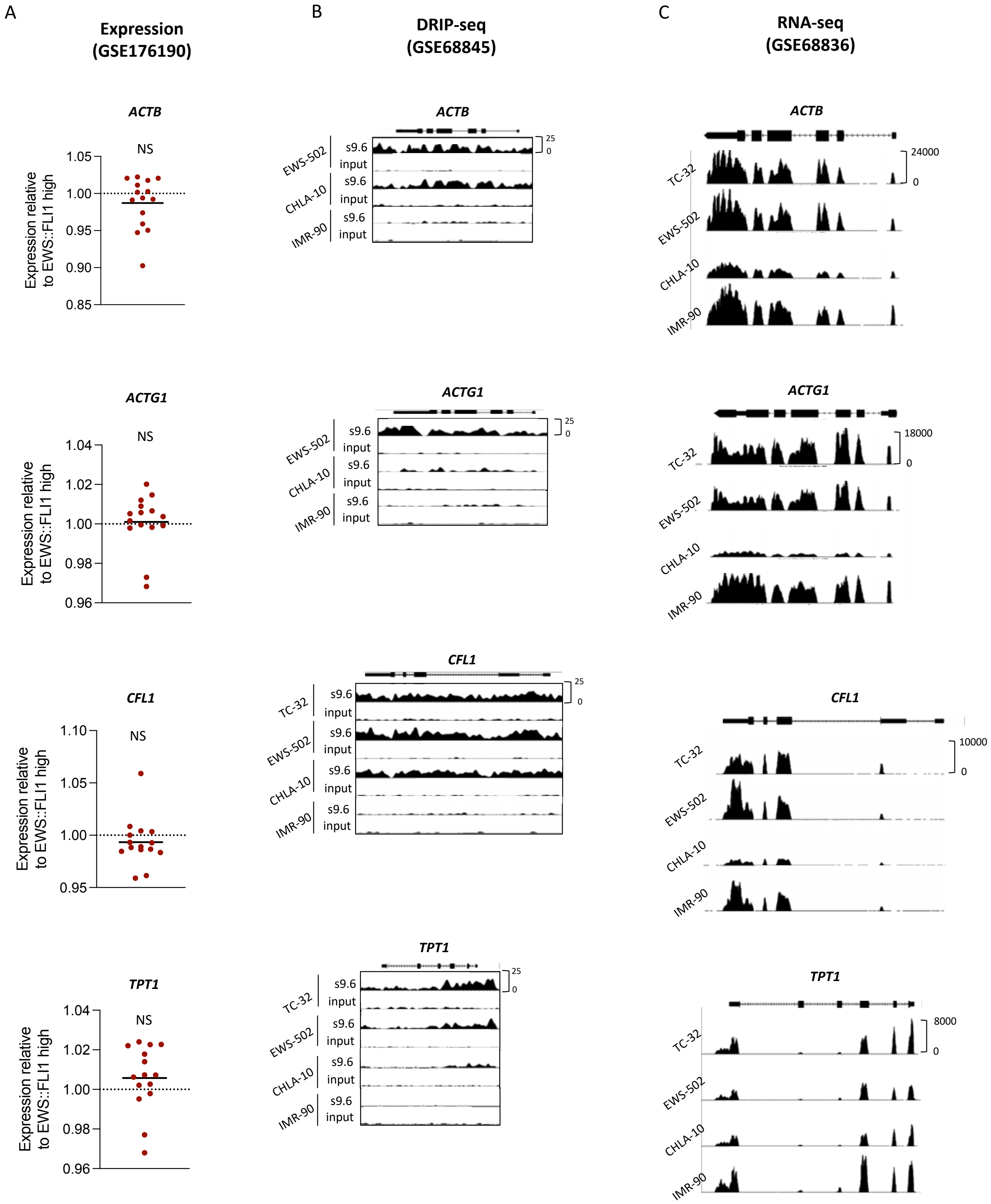

### Supplementary Figure 6.

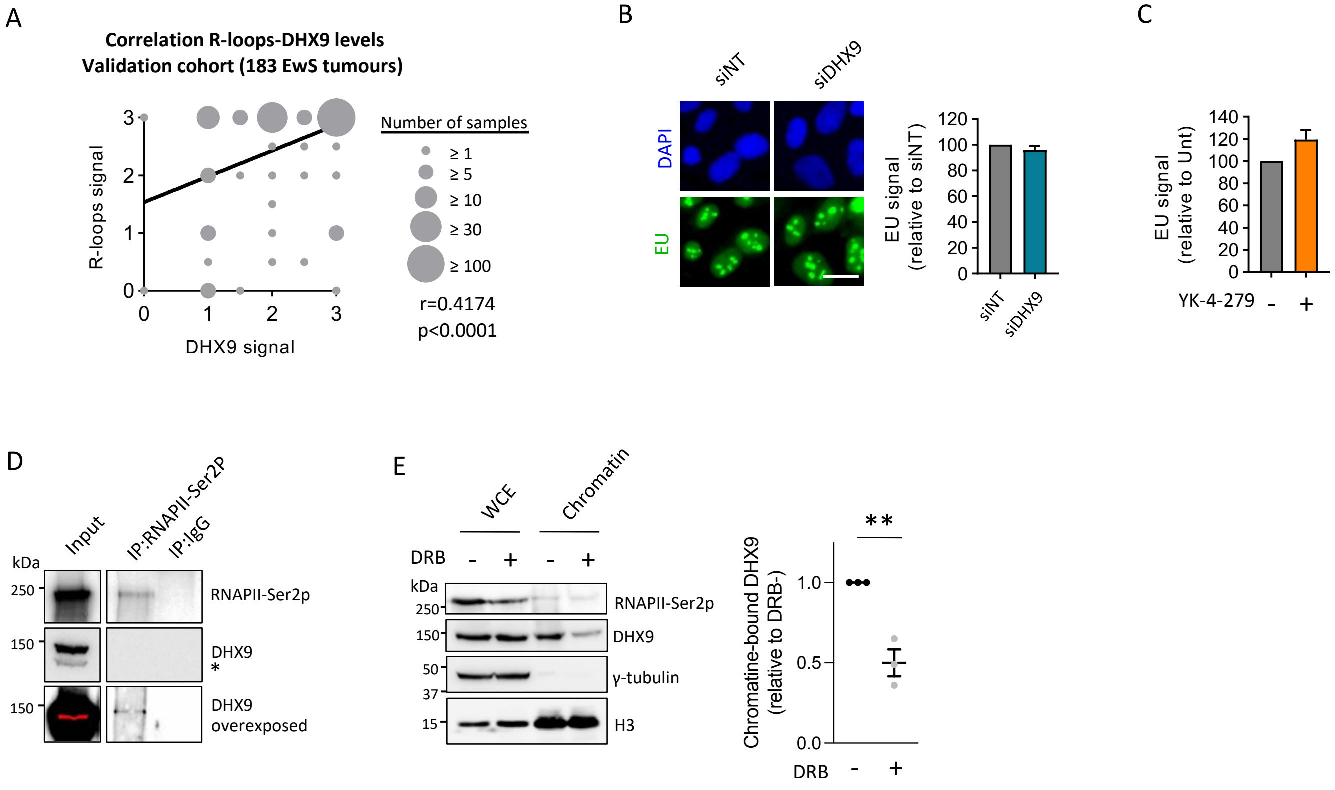
